# Peroxiredoxin 2 is required for the redox mediated adaptation to exercise

**DOI:** 10.1101/2022.12.14.520384

**Authors:** Qin Xia, José Carlos Casas-Martinez, Eduardo Zarzuela, Javier Muñoz, Antonio Miranda-Vizuete, Katarzyna Goljanek-Whysall, Brian McDonagh

## Abstract

Exercise generates a site-specific increase in Reactive Oxygen Species (ROS) within muscle required for a beneficial adaptive response by activation of specific signalling pathways. Here, we demonstrate that Peroxiredoxin 2 (Prdx2), an abundant cytoplasmic 2-Cys peroxiredoxin, is required for the adaptive beneficial hormesis response to H_2_O_2_. A short bolus addition of H_2_O_2_ increases mitochondrial capacity and improves myogenesis of cultured myoblasts, this beneficial adaptive response was suppressed in myoblasts with decreased expression of cytoplasmic Prdxs. A swimming exercise protocol in *C. elegans* increased mitochondrial content, fitness, survival and longevity in wild type (N2) worms. In contrast, *prdx-2* mutant worms had decreased fitness, disrupted mitochondria, reduced survival and lifespan following exercise. Global proteomics following exercise identified distinct changes in the proteome of N2 and *prdx-2* mutants. Furthermore, a redox proteomic approach to quantify reversible oxidation of individual Cysteine residues revealed a relatively more oxidised redox state following exercise in the *prdx-2* mutants. Our results demonstrate that conserved cytoplasmic 2-Cys Peroxiredoxins are required for the beneficial adaptive response to a physiological stress.

## Introduction

Exercise is essential to maintain and improve skeletal muscle function, it helps prevent age-related muscle atrophy and has systemic beneficial effects for all age-related diseases [1]. Exercise promotes mitochondrial biogenesis and induces changes in gene expression [2, 3]. During exercise there is a site specific increase in endogenous reactive oxygen species (ROS) within muscle fibres [4, 5], which is essential for the beneficial adaptive response to exercise and repair of muscle fibres [6, 7]. Skeletal muscle generates endogenous ROS during contractile activity primarily via members of NADPH oxidase (NOX) family located at the sarcolemma [4, 8]. Site specific exercise induced ROS generation is required for a range of metabolic and cellular events including mitochondrial biogenesis [9] and Glut4 translocation to the plasma membrane regulating glucose uptake during exercise [8]. Inhibiting ROS generation during exercise can blunt the beneficial adaptive responses to exercise and potentially the repair of muscle fibres [6, 7]. However, it remains to be understood how relatively non-specific oxidising molecules generated during contractile activity can have a specific signalling role, superoxide is generated extracellularly by NOX enzymes that is subsequently converted into H_2_O_2_ and transported into the muscle fibre.

Acute ROS generation directly and indirectly regulates the activity of key redox sensitive transcription factors involved in the antioxidant response Nuclear factor erythroid 2-like 2 (Nrf2), proteostasis via Forkhead (FOXO) family and inflammatory transcription factors such as Nuclear factor kappa B (NfkB) and Signal transducer and activator of transcription 3 (STAT3) [10-13], that are required for the adaptive response to exercise. Acute endogenous ROS generation and in particular H_2_O_2_, are signalling molecules that can directly affect the activity of regulatory proteins via reversible post-translational modifications of redox-sensitive Cysteine (Cys) residues. Due to the abundance and kinetics of Peroxiredoxins (Prdxs) reactivity with H_2_O_2_, it is highly unlikely that, at physiological concentrations of H_2_O_2_, any other proteins could compete with Prdxs for reaction with H_2_O_2_ [14]. Prdxs are peroxidases, however they have also been demonstrated to promote H_2_O_2_ induced oxidation of regulatory Cys residues on signalling proteins in yeast, mammalian cells and *C. elegans* models [12, 15-18].

The activation of exercise related transcription factors are either directly (STAT3, FOXO3) or indirectly (Nrf2, NfkB) regulated by the redox environment via oxidation of conserved Cys residues, that can fine tune the cellular response to acute and chronic stress. STAT3 has been identified as interacting with Prdx2 following a brief exposure to H_2_O_2_, forming a redox relay and attenuated transcriptional activity [12]. Similarly, FOXO members contain redox-sensitive Cys residues critical for function and likely involved in a redox signalling cascade regulating nuclear translocation [19]. Following endogenous H_2_O_2_ generation, such as during muscle contraction or injury, the flux of H_2_O_2_ through Prdxs located proximal to the site is predicted to outcompete alternative redox dependent signalling proteins [14, 20]. In contracted isolated single muscle fibres from adult mice, there is a rapid and reversible oxidation of Prdx2 that is blunted in muscle fibres from old mice [21]. Knockdown of Prdx2 in the mdx mouse model of muscle dystrophy where the expression of Prdx2 is significantly decreased compared to control mice, exacerbates the loss of muscle force production while overexpression of Prdx2 attenuates loss of muscle force [22]. In muscle biopsies from humans subjected to a high intensity interval training, a rapid reversible oxidation of mitochondrial Prdx3 has been demonstrated [23]. Mice deficient in Prdx3 do not show muscle atrophy but have a disrupted mitochondrial network, disrupted contractile function and faster rate of fatigue [24].

The nematode *Caenorhabditis elegans* is an excellent physiological model to dissect the mechanisms of adaptive signalling responses to exercise. The body wall muscle of *C. elegans* closely resembles the sarcomere of striated muscle in vertebrates and is a valuable model for investigation of muscle maintenance and function [25]. Similar to vertebrate species, body wall muscle from old worms undergoes nuclear fragmentation, disrupted mitochondrial dynamics, lipid accumulation and loss of muscle mass [26]. Moreover, a swim exercise regime has been demonstrated to improve mitochondrial capacity in body wall muscle and overall healthspan [27]. *C. elegans* do not possess orthologues of all the proteins or transcription factors that have been described in the mammalian response to exercise but contains 3 Prdx genes (*prdx-2, prdx-3* and *prdx-6*) and orthologues of Nrf2 (SKN-1), STAT (STA-1) and FOXO (DAF-16). It has previously been demonstrated that *prdx-2* mutants are short lived compared to wild types [28]. PRDX-2 is a typical cytoplasmic 2-Cys peroxiredoxin and has been demonstrated that the lifespan extension of *C. elegans* cultivated at lower temperatures and in response to metformin treatment requires the presence of PRDX-2 for mitohormesis and possible downstream activation of SKN-1 via p38 MAPK [29, 30]. Paradoxically PRDX-2 has also been identified as regulating insulin secretion and insulin dependent inhibition of SKN-1 [31]. In *C. elegans*, mitochondrial biogenesis and degradation are regulated by activation of SKN-1 [32]. Furthermore the mitophagy receptor protein BNIP3/Nix homologue in *C. elegans*, DCT-1, is among SKN-1 targets and its expression is non-redundantly co-regulated with DAF-16/FOXO transcription factor [32], all of which are linked to primary ROS generation and potentially a redox signalling cascade associated with PRDX-2. The generation of H_2_O_2_ by the activation of the dual oxidase BLI-3, can result in the downstream activation of SKN-1 via the p38 MAPK pathway but the precise mechanism responsible is unknown [33]. We hypothesise that PRDX-2 is an essential mediator for the adaptive hormesis response to endogenous ROS generation such as during exercise.

In this study, we demonstrate that a short bolus physiological concentration of H_2_O_2_, results in an exercise like adaptive response, an increase in mitochondrial capacity and improved myogenesis of cultured myoblasts. We demonstrate that Prdx2 is required for the beneficial adaptive response of myoblasts to H_2_O_2_ but is not required for myogenesis under normal conditions. Myoblasts with knockdown of Prdx1 and/or Prdx2 did not increase mitochondrial content following H_2_O_2_ treatment and had disrupted myogenesis. Following a 5-day *C. elegans* swimming protocol, previously demonstrated to improve healthspan, we observed increased mitochondrial content, improved fitness and survival in wild type worms [27]. Conversely, the *prdx-2* mutant worms subjected to the same exercise protocol had decreased fitness, disrupted mitochondria and reduced lifespan. A global proteomic approach revealed an increase in abundance of proteins involved in fatty acid oxidation and ion transport and a decrease in cuticle proteins following exercise regime in wild type N2 worms. An increase in abundance of proteins involved in mTOR pathway and actin binding proteins and a decrease in proteins involved in transsulfuration and microRNA biogenesis pathways were observed in the *prdx-2* mutant strain following exercise. Relative quantification of the reversible oxidation state of individual Cys residues revealed predominantly more relative oxidation of specific Cys residues following exercise in the *prdx-2* mutant. A number of specific regulatory Cys residues in metabolic, ribosomal and calcium handling proteins become more reduced in wild type worms but more oxidised in *prdx-2* mutants following exercise. Together, the data demonstrates the key role of Prdx2 as a regulator of intracellular redox signalling cascades required for effective adaptive hormesis response following endogenous ROS generation.

## Results

### Prdx1 and Prdx2 are sensors of physiological changes in H_2_O_2_

To determine the redox sensitivity of Prdxs in C2C12 skeletal muscle myoblasts treated with a range of H_2_O_2_ concentrations for 10 mins, samples were immediately homogenised in an alkylating buffer containing NEM to prevent thiol disulphide exchange. Following non-reducing Western blotting there was a significant shift in the dimer formation of Prdx2 at 10 μM of H_2_O_2_ and Prdx1 at 25 μM with no change in protein abundance and no effects on cell viability (Fig.1 a,b), there was no change in the monomer/dimer formation of Prdx3, 5 or 6 (Suppl. Fig1a-c). 10 min treatment at 50 μM (and above) of H_2_O_2_ resulted in the hyperoxidation of Prdxs (Fig.1c). A time course of Prdx oxidation following 25 μM H_2_O_2_ treatment resulted in significant changes in the dimer/monomer arrangement at 10 min and after 30 min there was a disappearance of hyperoxidised Prdxs with a return to the ratio of dimer/monomer formation as in controls (Suppl. Fig.1d-f), likely as a result of resolution of hyperoxidised Prdxs by sulfiredoxin or turnover of Prdxs. These results indicate that 10 min 25 μM H_2_O_2_ can induce reversible redox modifications of Prdx2.

**Figure 1.**
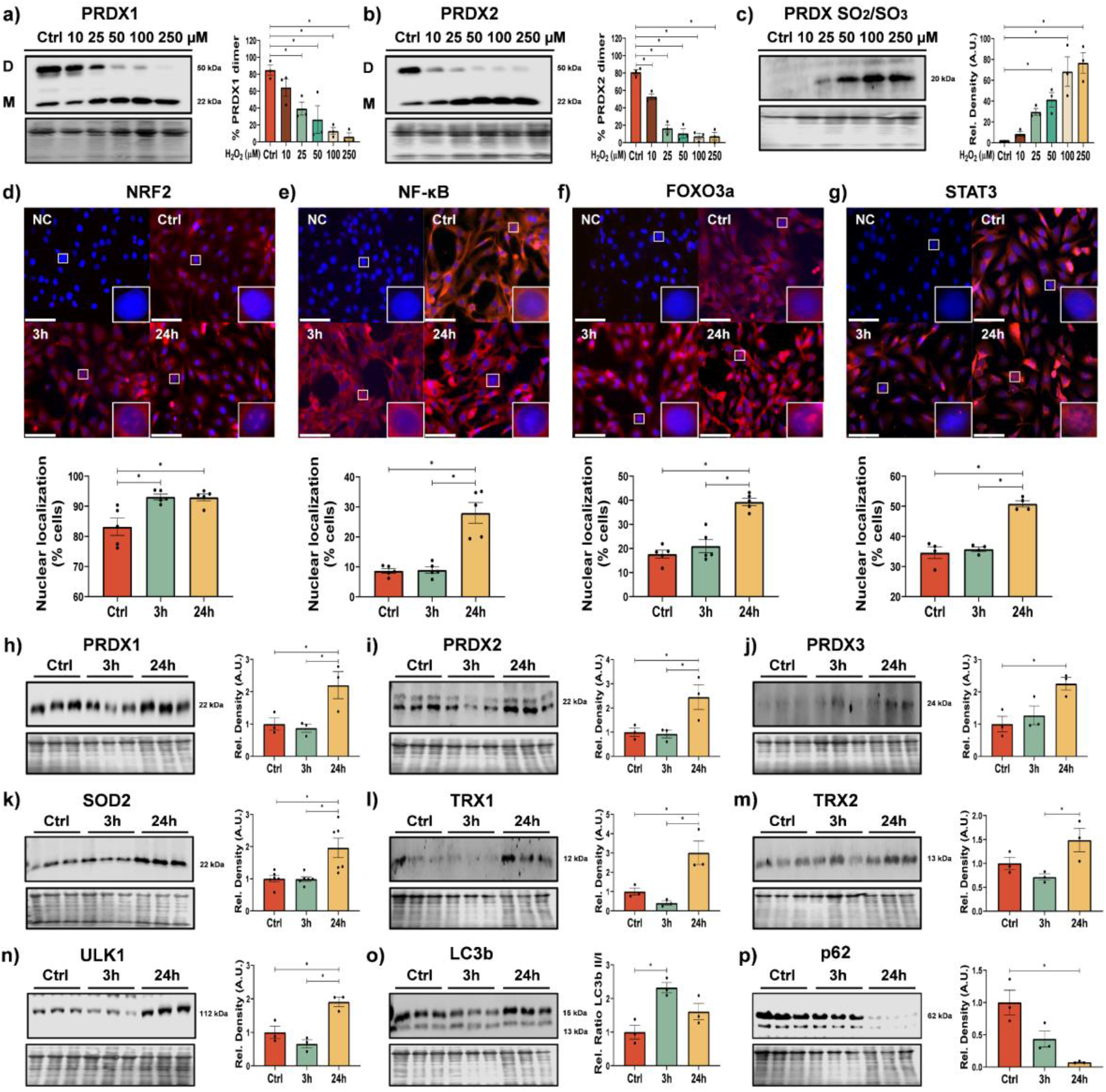
Acute treatment of H_2_O_2_ induces changes in redox state of Prdx1 and Prdx2 and increased nuclear localisation of redox sensitive transcription factors. Effects of different concentrations of H_2_O_2_ on monomer (M)/dimer (D) ratio of Prdx1 (a) and Prdx2 (b) and Prdx SO_2_/SO_3_ (c) in C2C12 myoblasts. Myoblasts were treated for 10 min with 25 μM H_2_O_2_, media was changed and % nuclear localisation of transcription factors NRF2 (d), NF-κB (e), FOXO3a (f) and STAT3 (g) was analysed after 3 and 24h; scale bar = 75 μm, n=3-5. Protein extracts and Western blotting was performed from either Ctrl or 3h and 24h following H_2_O_2_ treatment against PRDX1 (h), PRDX2 (i), PRDX3 (j), SOD2 (k), TRX1 (l), TRX2 (m), ULK1 (n) LC3B (o) and p62/SQSTM1 (p). Graphs are the normalised relative means +/-SEM and all experiments were performed with n=3 and *p-*value of <0.05 was considered as statistically significant *(*p*<0.05), one-way ANOVA was used for significance between groups (a-p). *p values (a: Ctrl vs 25μM= 0*.*0386; b: Ctrl vs 25μM< 0*.*0001; c: Ctrl vs 50μM= 0*.*0277; d: Ctrl vs 3h= 0*.*0066, Ctrl vs 24h= 0*.*0074; e: Ctrl vs 24h< 0*.*0001; f: Ctrl vs 24h< 0*.*0001; g: Ctrl vs 24h< 0*.*0001; h: Ctrl vs 24h= 0*.*0490; i: Ctrl vs 24h= 0*.*0468; j: Ctrl vs 24h= 0*.*0270; k: Ctrl vs 24h= 0*.*0067; l: Ctrl vs 24h= 0*.*0207; m: 3h vs 24h= 0*.*0355; n: Ctrl vs 24h= 0*.*0128; o: Ctrl vs 3h= 0*.*0091; p: Ctrl vs 24h= 0*.*0061)*.

### Adaptive response to physiological H_2_O_2_ results in improved mitochondrial capacity

Following skeletal muscle contraction, intracellular H_2_O_2_ increases to an estimated maximum of 50-100 nM [34]. The peroxide gradient across the cell membrane is estimated to be in the region ∼380-650 fold following a bolus addition of H_2_O_2_ to cells *in vitro*, depending on cell density and culture conditions [35, 36]. Accordingly, it is predicted that 25 μM of H_2_O_2_ would result in an intracellular concentration of ∼50 nM, similar to that obtained from contracting skeletal muscle during exercise. Following the bolus addition of H_2_O_2_ for 10 min, the media was replaced and cells were allowed to recover for 3h or 24h. There was a significant increase in nuclear localisation of exercise related transcription factors Nrf2, NF-kB, FOXO3a and STAT3 after 24h (Fig.1d-g). The abundance of Nrf2 target proteins such as Prdxs 1-3, SOD2 and Trx1 & 2, increased 24 h following 10 min bolus addition of H_2_O_2_ (Fig.1h-m). There was an increase in abundance of the autophagy regulator Ulk1, an increase in LC3II/I ratio indicating increased autophagy turnover (at 3h) and a decrease in autophagy adaptor protein p62/SQSTM1 abundance (Fig.1n-p). An increase in proteins related to mitochondrial turnover (DJ-1, Parkin and Bnip3) was also observed (Suppl. Fig.1g-i). This indicates that 24hr following a short bolus addition of H_2_O_2_ there is increased nuclear localisation of redox sensitive transcription factors, increased abundance of antioxidant proteins, increased expression of markers of autophagy and mitochondrial dynamics.

### Myoblasts treated with physiological levels of H_2_O_2_ result in enhanced myogenesis

As exercise induces an increase in mitochondrial content we investigated the effects of 10 min 25 μM H_2_O_2_ treatment on mitochondrial content and function in myoblasts. MitoTracker staining slightly increased in intensity at 24h following H_2_O_2_ treatment, however no change in mitochondrial ROS measured using MitoSOX was observed in H_2_O_2_ treated cells (Fig.2a). Seahorse analysis of respiration indicated an increase in basal and maximal respiration as well as an increase in spare respiratory capacity in myoblasts 24h following exposure to 10 min H_2_O_2_ compared to non-treated controls (Fig.2b). Proteins related to mitochondrial content (PGc1α, Tfam, Tom20) and STAT3 all increased in abundance (Fig.2c-f). These results indicate a short bolus addition of H_2_O_2_ to myoblasts generates a hormesis effect with increased mitochondrial capacity.

**Figure 2.**
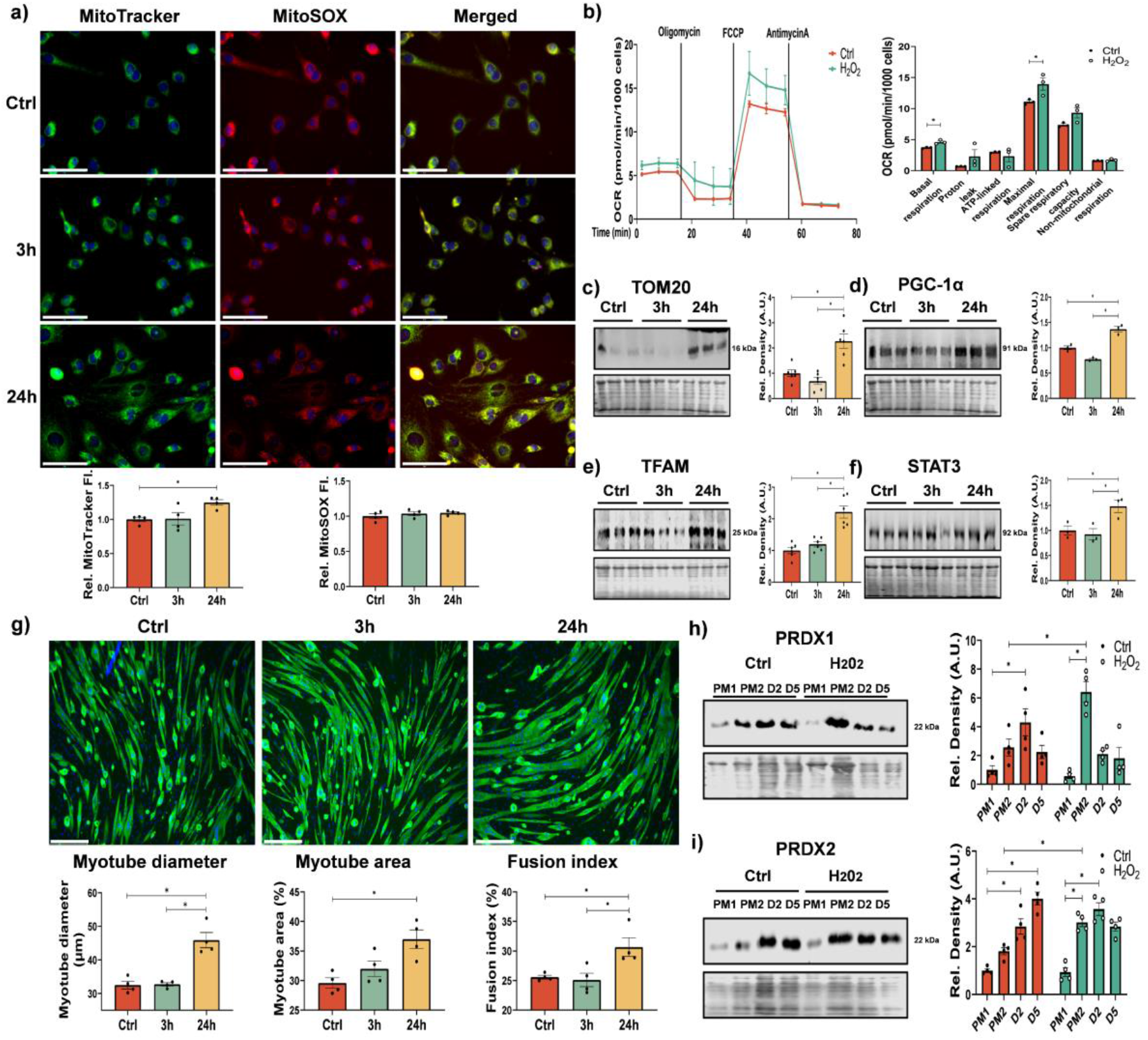
Hormesis effect of H_2_O_2_ improves mitochondrial capacity of myoblasts resulting in enhanced myogenic differentiation. Myoblasts treated for 10 min with 25 μM H_2_O_2_ were analysed for mitochondrial content using MitoTracker green and mitochondrial ROS using MitoSOX, n=4, one-way ANOVA used for significance between groups (a). Seahorse analysis of oxygen consumption of cells 24h following 10 min treatment with H_2_O_2_, n=3 and a Student *t* test used for analysis (b). Western blot analysis of proteins related to mitochondrial content TOM20 (c) and PGC1α (d) and TFAM (e) along with STAT3 abundance (f), n=3-6 one-way ANOVA used for significance between groups significance *p <0.05. MF 20 (antimyosin heavy chain; green) and DAPI (blue) immunostaining were performed for myogenic differentiation and nuclei identification, n=4 one-way ANOVA used for analysis, scale bar =75μm. (g). Protein extracts for Western blots of proliferating and differentiating myoblasts on different time points, growth media (GM), differentiation media (DM) Western blots of PRDX1 and PRDX2 expression at different time points, two-way ANOVA analysis was used for significance between groups (h,i). Graphs are the mean +/-SEM and all experiments were performed n=3-4 and *p-*value of <0.05 was considered as statistically significant *(*p*<0.05). *p values (a: MitoTracker fluorescence intensity Ctrl vs 24h= 0*.*0417, MitoSOX fluorescence intensity Ctrl vs 24h= 0*.*4744; b: Basal respiration: Ctrl vs H2O2= 0*.*0105, Maximal respiration: Ctrl vs H2O2= 0*.*0484, Spare respiration capacity: Ctrl vs H2O2= 0*.*0810; c: Ctrl vs 24h= 0*.*0010; d: Ctrl vs 24h= 0*.*0015; e: Ctrl vs 24h< 0*.*0001; f: Ctrl vs 24h= 0*.*0468; g: Myotube diameter: Ctrl vs 24h= 0*.*0004, Myotube area: Ctrl vs 24h= 0*.*0069, Fusion index: Ctrl vs 24h= 0*.*0273; h: Ctrl: PM1 vs Ctrl: D2= 0*.*0098, Ctrl: PM2 vs H2O2: PM2= 0*.*0019; i: Ctrl: PM1 vs Ctrl: D2< 0*.*0001, Ctrl: PM2 vs H2O2: PM2= 0*.*0127, H2O2: PM1 vs H2O2: PM2< 0*.*0001)*.

Myoblasts were exposed to H_2_O_2_ followed by a 24h recovery, the media was changed into differentiation medium and cells allowed to differentiate into myotubes. Myoblasts exposed to a short bolus concentration of 25 μM H_2_O_2_ were characterised by an increased myotube area, myotube diameter and fusion index, indicating improved myogenesis (Fig.2g). Expression levels of redox and mitochondrial proteins were analysed during the differentiation of myoblasts (Suppl Fig.2a-e and Fig.2h,i). Next, we examined whether the H_2_O_2_ induced changes in mitochondrial content were reflected in changes in Prdx levels. Interestingly, there was an initial increase in Prdx1 and subsequent decrease towards later stages of differentiation, while Prdx2 levels increased during differentiation. In turn, for H_2_O_2_ treated cells, Prdx1 and Prdx2 were initially expressed at higher levels 24h following treatment and subsequently the levels of Prdx1 declined and Prdx2 levels remained unchanged (Fig.2h,i). Expression levels of Trx2 and Parkin initially increased 24h following H2O2 treatment compared to non-treated cells while p62 decreased (Suppl Fig.2b-d). Results indicate that a short physiological addition of H_2_O_2_, similar to predicted values following contractile activity, improved mitochondrial capacity and enhanced myogenesis.

### Prdx1 and Prdx2 are required for myogenesis following a short bolus addition of H_2_O_2_ but not for myogenesis under normal conditions

In order to investigate the role of Prdx in myogenesis, we performed a transient knockdown of Prdx1 and/or Prdx2 using siRNA which we confirmed reduced protein levels to only ∼20% (Prdx1) and ∼40% (Prdx2) (Fig.3a,b). There was an increase in abundance of STAT3, TFAM, DJ-1 and BNIP3 in myoblasts initially treated with H_2_O_2_ (Fig.3c-f). Myoblasts increased SOD2 and decreased p62 expression following H_2_O_2_ treatment with knockdown of Prdx1 and/or Prdx2 along with increased Nrf2 nuclear localisation, as a response to increased oxidative stress (Suppl Fig.2f-h). The increase in the number of cells with nuclear localisation of STAT3 following H_2_O_2_ treatment was ablated when Prdx1 and/or Prdx2 were knocked down (Suppl Fig.2j). 24h following a bolus addition of H_2_O_2_ myoblasts were stained with MitoTracker Green and MitoSOX. As described in Fig. 2a, H_2_O_2_ treated myoblasts were characterised by increased mitochondrial content but no change in mitochondrial ROS were observed. However, myoblasts treated with Prdx1 and/or Prdx2 siRNA showed either similar (siPrdx1 and siPrdx2) or reduced (siPrdx1&2) mitochondrial content and an increase in MitoSOX staining suggesting an increase in mitochondrial ROS (Fig.3g). Myoblasts were subsequently differentiated into myotubes (Fig.3h). Myoblasts treated with siPrdx1 and/or siPrdx2, showed no change in myogenesis efficiency. However, myoblasts initially treated with 25 μM for 10 min H_2_O_2_ had improved myogenesis, whereas H_2_O_2_ treatment with knockdown of Prdx1 and Prdx2 abolished the adaptive response and enhanced myogenesis, instead these cells had decreased myotube diameter, area fraction and fusion index compared to controls, demonstrating disrupted myogenesis (Fig.3h). Myoblasts lacking either or both cytoplasmic 2-Cys Prdxs had dysregulated myogenesis and increased oxidative stress response as a result of loss of peroxidase activity. Together, the data demonstrate that Prdx1 and Prdx2 are essential for the beneficial adaptive hormesis response to a short bolus concentration of H_2_O_2_.

**Figure 3.**
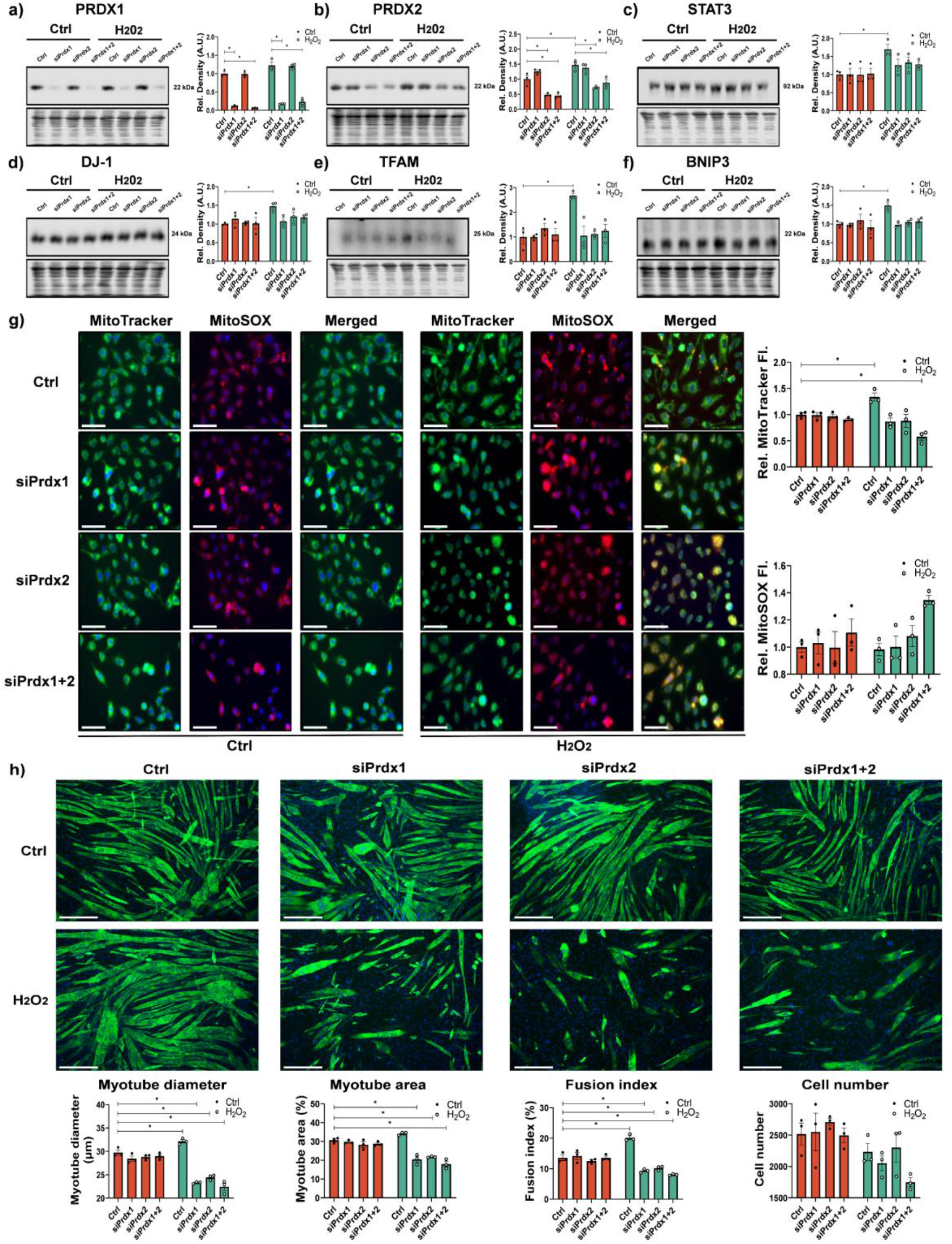
Knockdown of Prdx1 and/or Prdx2 inhibits hormesis effect of H_2_O_2_ and increases oxidative stress. Western blot confirming decreased expression PRDX1 and PRDX2 in C2C12 by siPrdx1 and/or siPrdx2 and 24h following 10 min 25 μM H_2_O_2_ treatment, n=3 (a,b). Increased expression of STAT3, Protein DJ-1, TFAM and BNIP3 24h following H_2_O_2_ treatment that was not detected with knockdown of Prdx1 and/or Prdx2 (c-f), n=3. MitoTracker green revealed increase in mitochondrial staining in H_2_O_2_ treated cells but not with knockdown of siPrdx1 and/or Prdx2, increase in MitoSOX staining following H_2_O_2_ treatment with siPrdx1+Prdx2 (g) n=3, scale bar =75μm. MF 20 immunostaining for myogenic differentiation revealed PRDX1 and PRDX2 not required for differentiation in non H_2_O_2_ treated cells but are required for hormesis effect and silencing results in disrupted myogenesis, scale bar =75μm (h). Graphs are the mean +/-SEM and all experiments were performed with n=3 and Two-way ANOVA analysis was used between groups and *p-*value of <0.05 was considered as statistically significant *(*p*<0.05) (a-h). *p values (a: Ctrl: Ctrl vs Ctrl: siPrdx1= 0*.*0176, Ctrl: Ctrl vs Ctrl: siPrdx1+2= 0*.*0110, H2O2: Ctrl vs H2O2: siPrdx1< 0*.*0001, H2O2: Ctrl vs H2O2: siPrdx1+2< 0*.*0001; b: Ctrl: Ctrl vs Ctrl: siPrdx2= 0*.*0078, Ctrl: Ctrl vs Ctrl: siPrdx1+2= 0*.*0050, Ctrl: Ctrl vs H2O2: Ctrl= 0*.*0168, H2O2: Ctrl vs H2O2: siPrdx2< 0*.*0001, H2O2: Ctrl vs H2O2: siPrdx1+2< 0*.*0001; c: Ctrl: Ctrl vs H2O2: Ctrl= 0*.*0468; d: Ctrl: Ctrl vs H2O2: Ctrl= 0*.*0420; e: Ctrl: Ctrl vs H2O2: Ctrl= 0*.*0018; f: Ctrl: Ctrl vs H2O2: Ctrl= 0*.*0472; g: MitoTracker fluorescence intensity: Ctrl: Ctrl vs H2O2: Ctrl= 0*.*0445, Ctrl: Ctrl vs H2O2: siPrdx1+2= 0*.*0083; h: Myotube diameter: Ctrl: Ctrl vs H2O2: Ctrl= 0*.*0414, Fusion index: Ctrl: Ctrl vs H2O2: Ctrl< 0*.*0001)*.

### Improved mitochondrial capacity and survival following swimming exercise protocol in C. elegans that is suppressed in prdx-2 mutants

In order to investigate the adaptive response to exercise in the context of a complete organism we moved to *C. elegans* wild type and mutant strains. We performed a 2×90 min daily swimming protocol using *C. elegans* for 5 days that has previously been demonstrated to improve healthspan, locomotory function and inhibit deterioration in models of neurodegenerative disease [27]. We used *C. elegans* N2 wild type strain and mutants *prdx-2 (gk169), skn-1 (zj15)* and *bli-3 (e767)*, which encode key enzymes involved in redox homeostasis. The *prdx-2(gk169)* mutation is a deletion strain with a presumably null allele for *prdx-2*, the single homologue of human Prdx1 and Prdx2. SKN-1 is the *C. elegans* functional orthologue of mammalian Nrf2 transcription factor, the *skn-1(zj15)* strain has disrupted splicing and reduced *skn-1* mRNA. The dual oxidase *bli-3* is one of two dual oxidase (duox) NOX related genes in *C. elegans* (duox1/bli-3 and duox2), which share 94% amino acid similarity but the function of duox2 is unknown [37]. The *bli-3(e767)* strain contains a mutation in the peroxidase domain of *bli-3* generating a reduction of function allele. As previously reported the *prdx-2, skn-1* and *bli-3* mutant strains had reduced longevity compared to the wild type N2 strain (Fig.4a) [28, 38, 39]. BLI-3 is required for pathogen resistance and promotes stress resistance via redox signalling cascade via activation of SKN-1 [38]. However, the *bli-3 (e767)* strain were very fragile, with a poor ability to survive following the exercise protocol and were therefore not used further. The 5day swimming protocol outlined in Fig.4b, improved lifespan of N2 worms but decreased lifespan of *prdx-2* mutants and had no effect on lifespan of *skn-1* mutants (Fig.4c). Moreover, the swimming protocol increased survival of N2 worms, when exposed to the mitochondrial redox cycler Paraquat or peroxide treatment with tert-butyl hydroperoxide (TBH) (Fig.4d,e). However, in *prdx-2* mutants the swimming protocol resulted in increased sensitivity to both Paraquat and TBH (Fig.4d,e). In contrast, there was no change in the survival of *skn-1* mutants to TBH or Paraquat following exercise. These results highlight the requirement of PRDX-2 for the beneficial adaptation to exercise, and detrimental effect of exercise in strains lacking PRDX-2.

**Figure 4.**
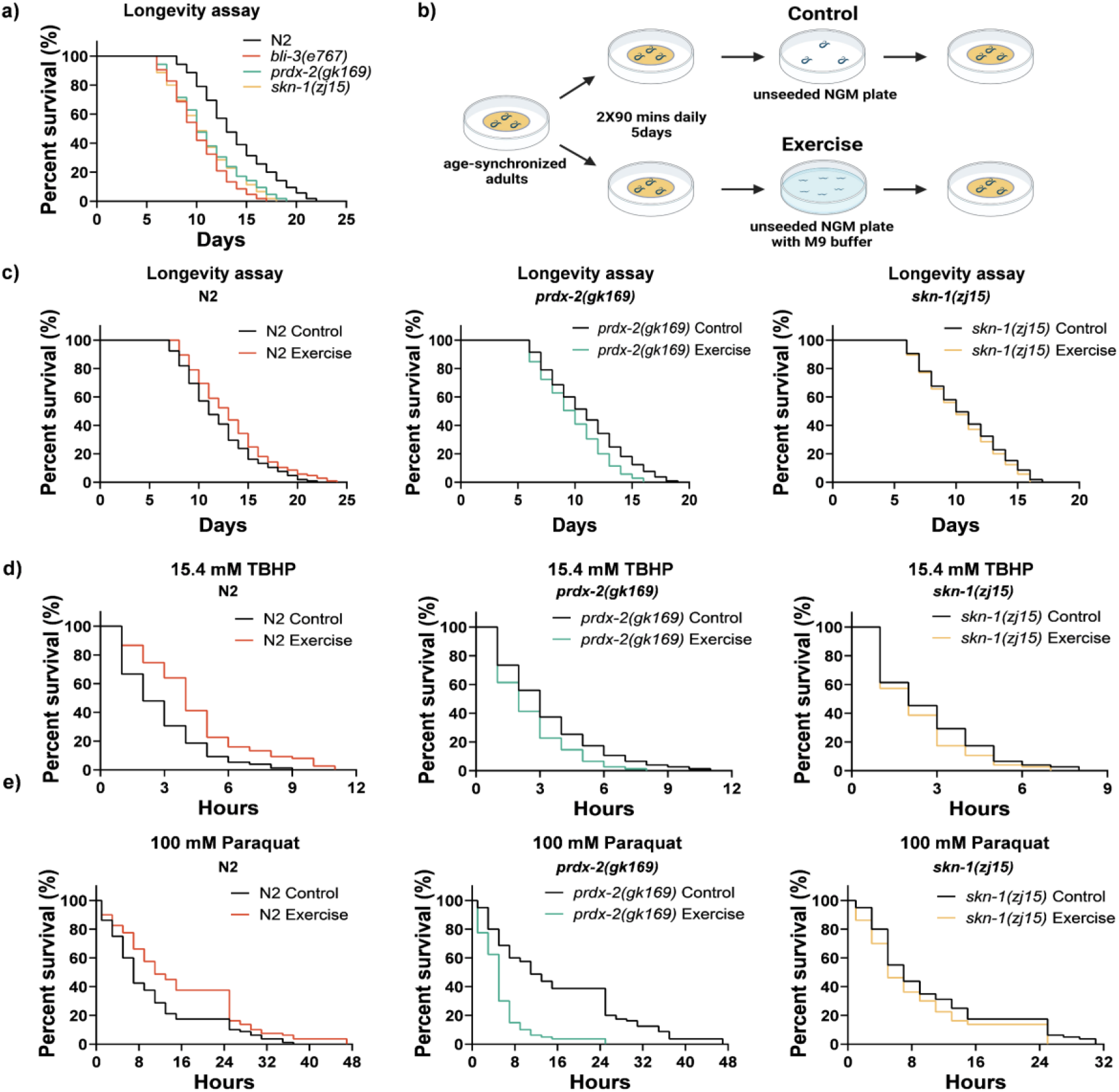
PRDX-2 is required for the improved survival and longevity following 5day exercise protocol. Longevity assays of *C. elegans* strains N2, *prdx-2 (gk163), skn-1 (zj15)* and *bli-3 (e767)*. Decreased longevity of *prdx-2, skn-1* and *bli-3* strains compared to N2 wild types (a). Schematic of swimming protocol for C. elegans strains (b). Following 5d exercise protocol worms were seeded on NGM plates and lifespan recorded, increased lifespan in exercised N2 worms, decreased lifespan in *prdx-2* strains and no change in *skn-1* mutants following exercise (c). Increased survival of N2 strain following exercise protocol when exposed to TBHP or paraquat, decreased survival of *prdx-2* mutants and no change in *skn-1* mutants following 5 day exercise protocol (d-e). Longevity experiments used a minimum of 105 worms, survival assays used minimum of 80 worms per condition. Log-rank (Mantel-Cox) analysis was used to evaluate survival between groups. *p values (a: N2 vs bli-3 (e767)< 0*.*0001, N2 vs prdx-2 (gk163)< 0*.*0001, N2 vs skn-1 (zj15) < 0*.*0001; c: N2: Control vs Exercise= 0*.*0266; prdx-2 (gk163): Control vs Exercise= 0*.*0023; skn-1 (zj15): Control vs Exercise= 0*.*4060; d: N2: Control vs Exercise< 0*.*0001; prdx-2 (gk163): Control vs Exercise= 0*.*0112; skn-1 (zj15): Control vs Exercise= 0*.*1592; e: N2: Control vs Exercise= 0*.*0033; prdx-2 (gk163): Control vs Exercise< 0*.*0001; skn-1 (zj15): Control vs Exercise= 0*.*0886)*.

Using the zcIs14 [*myo-3p::mitogfp*] muscle mitochondrial reporter strain, we observed that the swimming protocol resulted in increased filamentous mitochondria and decreased mitochondrial punctae in body wall muscle (Fig5a). The exercise protocol also resulted in increased mitochondrial turnover as assessed by the mitophagy reporter strain [*myo-3p tomm-20::Rosella*] (Fig. 5b). The Rosella biosensor contains a pH-stable red fluorescent protein (RFP) fused to a pH-sensitive green fluorescent protein (GFP), the mitochondrial network fluoresces red and green under normal conditions. During mitophagy, with increased delivery of mitochondria to the acidic environment of lysosomes, GFP fluorescence is quenched while RFP remains stable [40]. Basal and maximal respiration were increased in N2 wild type worms subjected to swimming protocol compared to non-exercised controls an effect not detected in *prdx-2* or *skn-1* mutants (Fig.5c). Furthermore, the swimming assay, resulted in increased SKN-1 activity as measured by fluorescence using the *skn-1* activation reporter strain *gst-4p::gfp* (Fig.5d). MitoTracker Red staining intensity increased in N2 worms following swimming, suggesting increased mitochondrial membrane potential. This increase was not observed in the *skn-1* mutants and *prdx-2* mutants had decreased staining intensity following the swimming protocol (Fig.5e). Mitochondrial ROS was assessed using MitoSOX. No difference in mitochondrial ROS was detected following the swimming protocol in N2 or *skn-1* strains, whereas there was a significant increase in ROS following exercise in the *prdx-2* mutants (Fig.5f). Using reducing and non-reducing gels, we assessed the expression and dimer formation of PRDX-2 following exercise in the N2 and *skn-1* strains. There was no change in PRDX-2 protein abundance but a shift in dimer/monomer formation in N2 worms and a slight increase in abundance of PRDX-2 in *skn-1* strain following exercise, no PRDX-2 was detected in *prdx-2* mutants (Suppl Fig.3 a-d). Body size differences between the different strains (Suppl Fig.3e) make it difficult to compare the direct effects of exercise across strains (e.g. oxygen consumption), we therefore compared responses to exercise to non-exercised controls in each strain. The 5-day swimming protocol in N2 worms improved lifespan, increased their ability to withstand stress, increased mitochondrial content and respiration. These adaptive responses were repressed in *prdx-2* and *skn-1* mutants. Furthermore, the swimming exercise led to the *prdx-2* mutant being more susceptible to stress and decreased lifespan following exercise. Exercise had no effects on the survival assay of *skn-1* mutants. This suggests that while SKN-1 is activated during exercise in N2 strains, consistent with its activation following a physiologically relevant redox stress, this adaptation to exercise induced redox stress is mediated via PRDX-2.

**Figure 5.**
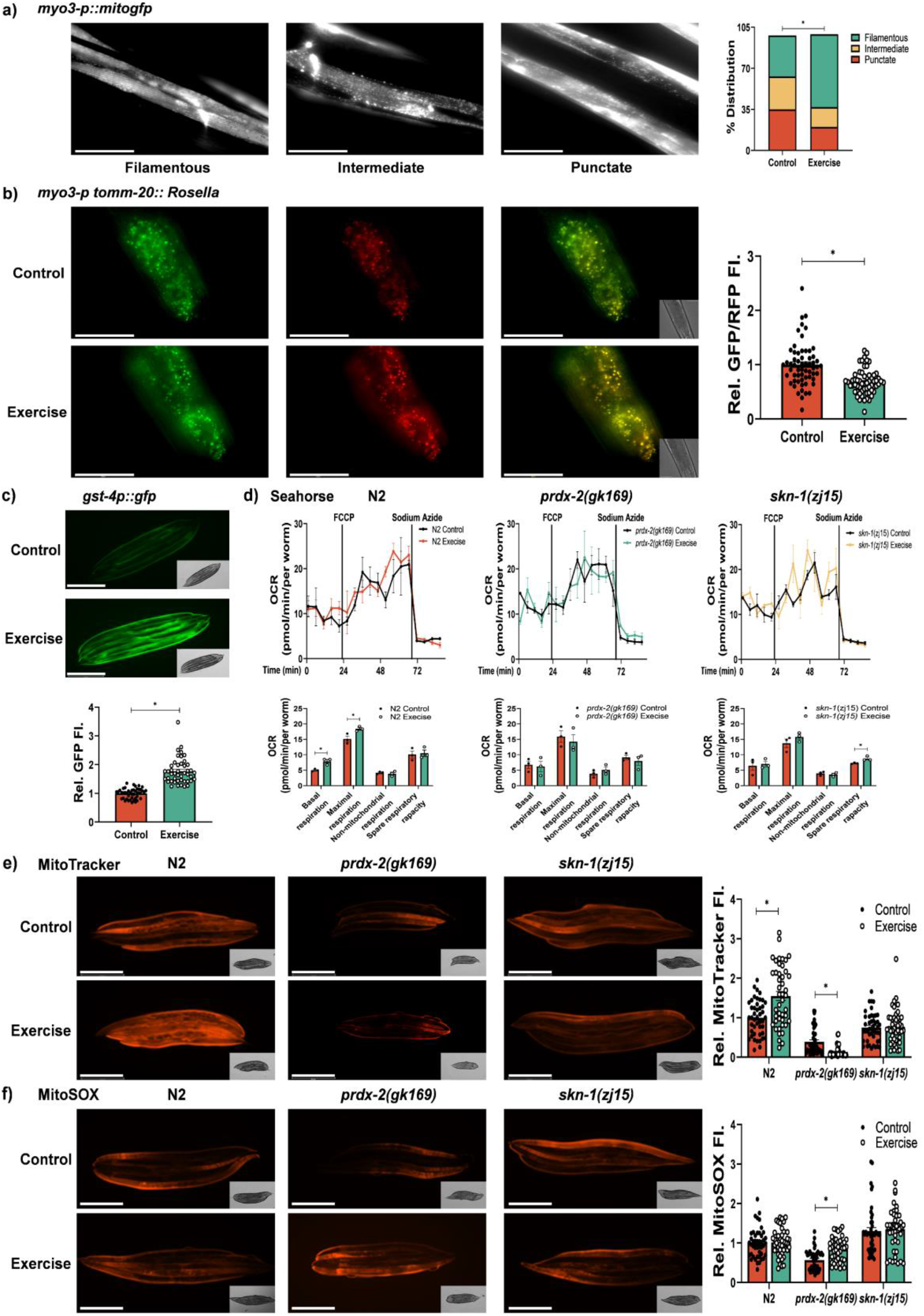
Exercise protocol promotes mitochondrial dynamics, function and content but not in prdx-2 mutants. Increase in filamentous muscle mitochondria following 5day exercise protocol using the *myo-3::gfp* reporter strain and scored according to [41], scale bar = 50μm (a). Representative images of *myo3-p::tomm20::Rosella* mitophagy reporter following 5 day swimming protocol (b). Representative images of *gst-4p::gfp* SKN-1 transcriptional reporter following 5day exercise protocol, data is represented as fluorescence intensity and experiments performed at least 3 times, scale bar = 275μm with at least 45 worms per strain/experiment (c). Seahorse analysis of oxygen consumption at basal, maximal (following FCCP) and non-mitochondrial (following sodium azide) respiration in non-exercised and exercised worms (d). Experiments were performed in triplicate with 8-13 worms per replicate. MitoTracker red staining of worms for mitochondrial potential (e) and MitoSOX staining for mitochondrial ROS (e) in non-exercised and exercised worms, scale bar=275μm. Graphs are the mean +/-SEM and all experiments were repeated at least 3 times and *p-*value of <0.05 was considered as statistically significant *(*p*<0.05). *p values (a: Control vs Exercise= 0*.*0008; b: Control vs Exercise< 0*.*0001; c: Control vs Exercise< 0*.*0001; d: Basal respiration: N2 Control vs Exercise= 0*.*0112, Maximal respiration: N2 Control vs Exercise= 0*.*0318; e: N2: Control vs Exercise< 0*.*0001, prdx-2 (gk163): Control vs Exercise< 0*.*0001; f: prdx-2 (gk163): Control vs Exercise< 0*.*0001)*.

### Global and redox proteomics following exercise in wild type and mutant C. elegans prdx-2 and skn-1

In order to determine the mechanisms underlying adaptation to exercise and mediating the phenotypes of the mutant worms, we analysed the changes in the proteome induced by the 5d exercise protocol in wild type and mutant strains through shotgun and redox proteomics. Exercise generates endogenous ROS that can be relayed during signal transduction via the reversible redox modifications of sensitive Cys residues. The proteomic approach included a differential Cys labelling step of reduced (NEM, light) and reversibly oxidised (D5 NEM, heavy) Cys residues to relatively quantify the reversible oxidation of individual Cys residues from redox sensitive residues using the ratio of D5 NEM:NEM (Suppl Fig4. Schematic of redox proteomic approach) [42]. Using this approach 3,590 proteins were identified and quantified for the global proteomic approach and 1,057 Cys containing redox peptides labelled with both NEM and D5 NEM were quantified for the redox proteomic analysis (Suppl. Files 1,2). Exercise can induce large scale transcriptional and proteomic changes. Distinct changes in the proteome of *prdx-2* and *skn-1* mutant strains were detected in response to the swimming protocol compared to wild type worms. We considered proteins that had an adjusted p-value <0.01 and at least a fold change > 1.5 in abundance as significantly changed as a result of exercise in each strain (Fig.6a-c).

**Figure 6.**
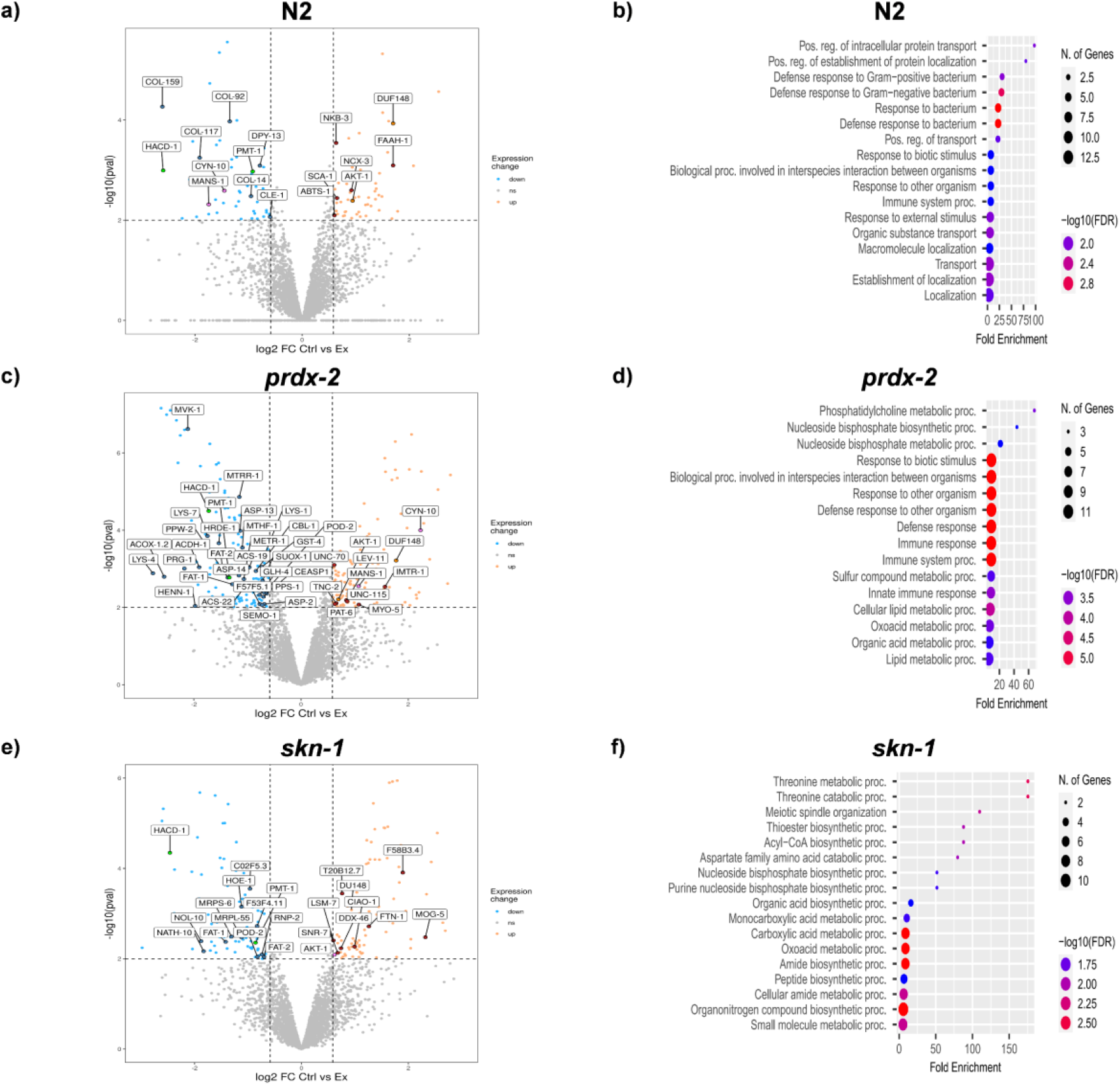
Global proteomics identifies distinct proteomic signatures following exercise and redox proteomics reveals Cys oxidation following exercise in prdx-2 mutants. LFQ proteomic data represented by volcano plots for changes in protein abundance following exercise, N2 (FDR=5.14%) (a) and ShinyGO enrichment of pathways of significant upregulated proteins N2 (b). Volcano plot of LFQ data in *prdx-2* mutant (FDR=3.8%) (c) and Shiny GO enrichment of significantly downregulated proteins following exercise (d). Volcano plot of LFQ data in *skn-1* mutant (FDR=6.5%) (e) and ShinyGO enrichment of downregulated proteins following exercise (f). Proteomics was performed from 4 biological replicates per condition and with ∼ 100 worms per replicate.

Exercise resulted in an increase in abundance of 66 proteins and a decrease in 67 proteins in N2 worms compared to non-exercised controls (Fig.6a). Proteins involved in metabolic function such as worm orthologues of serine/threonine protein kinase (AKT-1) proteins involved in fatty acid beta oxidation (FAAH-1) and proteins involved in Na/K and calcium transport (NKB-3, SCA-1, NCX-3 and ABTS-1) were increased with exercise compared to controls. Downregulated proteins included proteins involved in fatty acid anabolism (HACD-1) and proteins involved in cuticle formation (CLE-1, COL-14, −92, −117, −159, −166, −179 and DPY-13). ShinyGO analysis of N2 mutants highlighted the enrichment of intracellular protein transport as upregulated proteins and anatomical structural development proteins as downregulated (Fig.6d). In the *prdx-2* mutant, exercise resulted in an increase in abundance of 100 proteins and a decrease in abundance of 112 proteins and enrichment (Fig.6c). Proteins involved in mTOR activation (AKT-1, DAF-18, GSK-3, RHO-1 and R148.2) and actin binding proteins (LEV-11, MYO-5, PAT-6, UNC-70 and UNC-115) and troponin C (TNC-2), were all increased in exercised *prdx-2* worms. Proteins that decreased in *prdx-2* mutants compared to non-exercised controls included proteins involved in fatty acid synthesis (FAT-1, -2, ACDH-1, ACS-22, HACD-1, POD-2 and F08A8.2), transsulfuration pathways (CBL-1, METR-1, MTHF-1, MTRR-1, PPS-1, SEMO-1, SUOX-1, ACS-19, MVK-1 and GST-4) proteins involved in miRNA biogenesis (HRDE-1, PPW-2, HENN-1, PRG-1 and GLH-4) and peptidases (ASP-2, ASP-13, ASP-14, CEASP1, LYS-1, LYS-4, LYS-7 and F57F5,1) (Fig.6c). ShinyGO identified an enrichment of phosphatidylcholine and nucleoside bisphosphate metabolism in significantly downregulated proteins (Fig.6d). In the *skn-1* mutants the abundance of 54 proteins increased and 61 decreased in abundance following exercise (Fig.6c). Proteins that increased in abundance include those involved in iron metabolism (T20B12.7, CIAO-1, Y18D10A.9 and FTN-1) and RNA binding (LSM-7, SNR-7, MOG-5, DDX-46 (Protein F53H1.1) and F58B3.4. Proteins that decreased include the fatty acid synthesis proteins (FAT-1, FAT-2, HACD-1 and POD-2) and ribonucleoproteins (CO2F5.3, F53F4.11, HOE-1, MRPL-55, MRPS-2, MRPS-6, NATH-10, NOL-10, and RNP-2) (Fig.6c). ShinyGO identified an enrichment of threonine and acyl-CoA metabolism in significantly downregulated proteins (Fig6e).

Proteins commonly regulated in all 3 strains following exercise include the key metabolic protein AKT-1 and DUF148 domain containing protein, which increased in abundance following exercise. Proteins that decreased in abundance following exercise in all three stains were HACD-1 (Hydroxy-Acyl-CoA Dehydrogenase) predicted to be involved in beta oxidation of fatty acids and PMT-1 (Phosphoethanolamine N-methyltransferase 1) which catalyses the first step in the synthesis of phosphocholine and COL-166 (structural constituent of cuticle). Two proteins decreased in abundance in the N2 strain but increased in the *prdx-2* mutant strain following exercise, CYN-10 (Peptidyl-prolyl cis-trans isomerase-like 3) and MANS-1 (alpha-1,2-Mannosidase). Supplementary file 1 contains the full list of quantified proteins in each strain. Overall, the global proteomic approach generated a very distinct proteomic profile from each of the strains in response to exercise, the increase in abundance of ion regulatory proteins and decrease in cuticle proteins in the N2 strain. In the *prdx-2* mutant strain an increase in abundance of proteins in the mTOR pathway and proteins involved in the regulation of the cytoskeleton and a decrease in proteins involved in transsulfuration, peptidases and miRNA biosynthesis were detected following exercise.

The redox proteomic approach labelled reduced Cys residues with the thiol alkylating reagent NEM, followed by desalting of samples to remove excess NEM and reduction of reversibly oxidised Cys residues, the newly reduced Cys were labelled with D5 NEM (Suppl Fig.4). The ratio of the Cys containing peptide intensity labelled with both NEM and D5 NEM, allows the relative quantification of the reversible oxidation state of individual Cys containing peptides and the ratios of individual Cys residues from non-exercised and exercised strains were compared following exercise. Overall 1,057 Cys containing peptides labelled with both NEM and D5 NEM were quantified. We considered a p-value of < 0.05 and a log_2_FC > 1.5 (fold change 2.8) in the ratio of D5 NEM/NEM labelling following exercise protocol as significantly changed (Suppl file 2). Supplementary Table 1 is list of Cys containing peptides of where the redox state is significantly changed comparing the Log_2_Fold change of ratio of Heavy NEM:NEM labelling of Cys residues in non-exercised strains N2 Vs *prdx-2* and N2 Vs *skn-1* and following exercise compared to non-exercised controls in N2, *prdx-2* and *skn-1* strains.

Comparing the redox state of Cys residues in non-exercised N2 compared to both mutant *prdx-2* and *skn-1* non-exercised strains, the overall redox state of Cys residues are more reduced in both the *prdx-2* and *skn-1* strains (Fig.7a,b). In the wild type N2 strain, exercise induced changes in the redox state of Cys residues with specific Cys residues becoming both more reduced and others more oxidised (Fig.7c). However, mutants lacking functional PRDX-2 had a relatively more oxidised redox state following exercise with a higher ratio of D5-NEM:NEM (shift to right in volcano plot relatively more heavy NEM bound) (Fig.7d). Similarly, in the *skn-1* mutant following exercise more Cys residues become oxidised (Fig.7e). Fig.7f includes representative XICs of RPS-28_Cys32 and CRT-1_Cys133 in the different strains with and without exercise.

**Figure 7.**
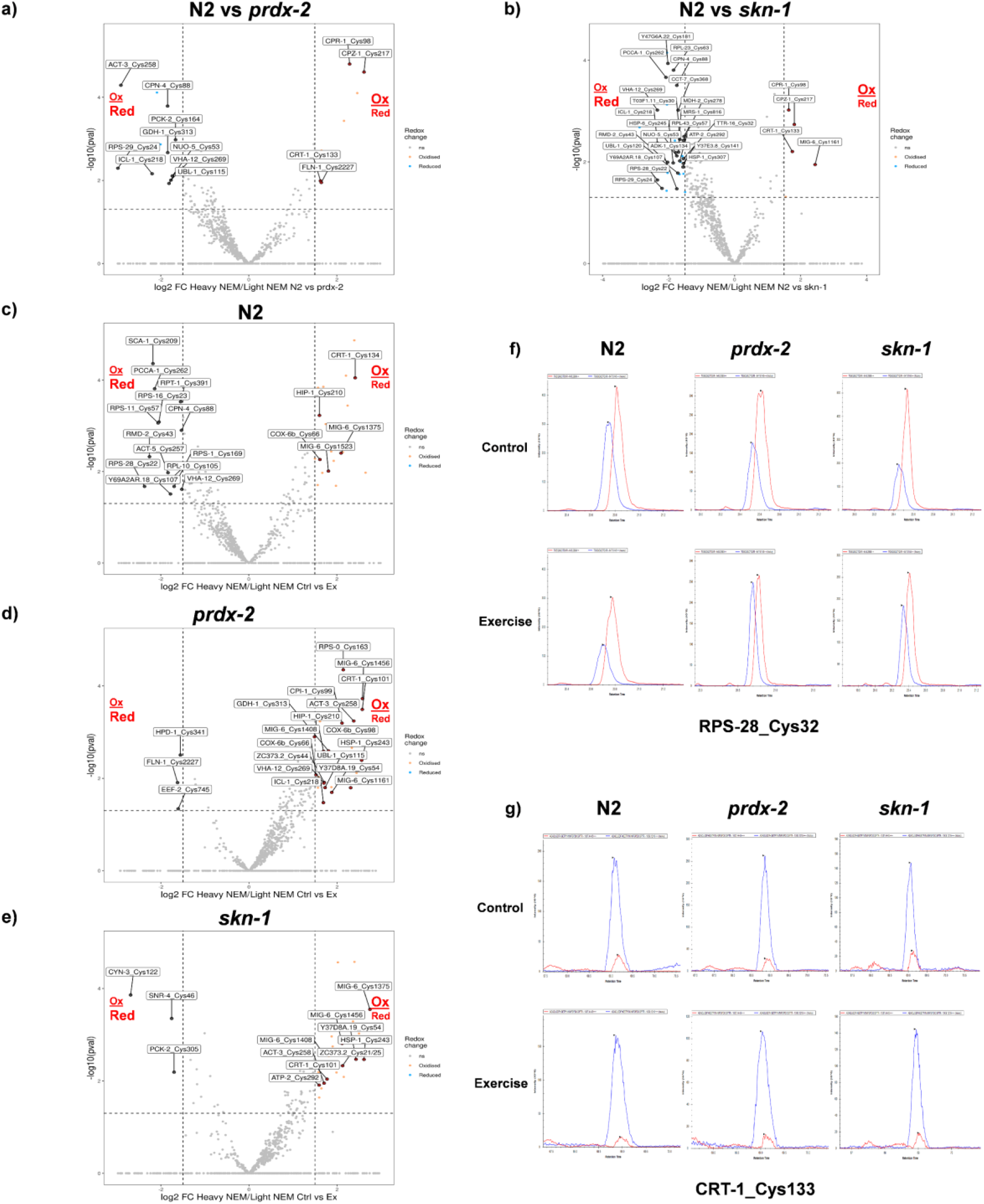
The prdx-2 mutant Cys are more reduced compared to N2 strain in non-exercised worms but are more oxidised following exercise. Volcano plots of redox proteomic data Log_2_Fold change of ratio Heavy:Light NEM labelled Cys containing peptides. Cys containing peptides that are relatively more oxidised following exercise shift towards the right and Cys residues that are more reduced following exercise shift towards the left. Non-exercised N2 Vs non-exercised *prdx-2* mutant (FDR=10.28%) (a) N2 Vs non-exercised *skn-1* mutant (FDR=9.67%) (b), N2 Ctrl Vs Exercised (FDR=8.79%) (c), *prdx-2* Ctrl Vs Exercised (FDR=12.75%) (d) and *skn-1* Ctrl Vs Exercised (FDR=10.07%) (e). Representative XICs of selected redox sensitive Cys residues from RPS-28 Cys32 (f) and CRT-1 Cys132 (g) from non-exercised and exercised samples. Peptides with reduced Cys residues are labelled with light NEM and represented in red, peptides with reversibly oxidised Cys residues are labelled with heavy NEM and represented in blue (f). Proteomics was performed from 4 biological replicates per condition and with ∼ 100 worms per replicate.

In the wild type N2 strain, 22 Cys residues from 21 proteins had a significant more relatively oxidised state and 13 Cys residues more reduced state following the exercise protocol (Fig.7b). In the *prdx-2* mutant strains, a very distinct profile in the redox state of Cys residues was observed, with 24 Cys resides from proteins becoming more oxidised and 3 Cys residues more reduced following exercise (Fig.7c). In the *skn-1* mutant, there were 23 Cys residues from proteins more oxidised and only 3 Cys residues more reduced following exercise (Fig.7d). We focused on comparing the shift in redox specific changes on the sensitive Cys residues across the different strains in response to exercise. If PRDX-2 is involved in a redox signalling cascade involving the transfer oxidising equivalents to target proteins it would be expected that Cys residues have a relatively more reduced state in non-exercised *prdx-2* mutants compared to N2 strain (Fig.7a). Cys containing proteins identified as more reduced in response to exercise in the wild type but are more oxidised or have no change following exercise in the *prdx-2* mutant strain could be potential targets of PRDX-2 in a redox relay and required for the adaptive response to exercise. In N2 wild type strain only, a number of proteins have Cys residues that are more reduced following exercise including components of the ribosome (RPS-1_Cys169, RPS-11_Cys57, RPS-16_Cys23, RPS-28_Cys32 and RPL-10_Cys105), mitochondrial proteins and regulatory proteins including ACT-5_Cys257 (Actin), CPN-4_Cys88 (Calponin), PCCA-1_Cys262 (Propionyl-CoA carboxylase alpha chain, RMD-2_Cys43 (TPR_REGION domain-containing protein), RPT-1_Cys391 (26S proteasome regulatory subunit 7), SCA-1_Cys209 (Calcium-transporting ATPase), Y69A2AR.18_Cys107 (Proton transporting ATP synthase) and VHA-12_Cys269 (V-type proton ATPase subunit B 1). In particular Cys269 from VHA-12 although more reduced in the N2 strain, it was more oxidised in the *prdx-2* strain following exercise. In the *prdx-2* mutant 3 Cys residues become more reduced following exercise Calponin (FLN-1_Cys2227), elongation factor 2 (EEF-2_Cys745) and 4-hydroxyphenylpyruvate dehydrogenase (HPD-1_Cys341). In the *skn-1* mutant Peptidyl-prolyl isomerase 3 (CYN-3_Cys122), small nuclear ribonucleoprotein (SNR-4_Cys46) and Phosphoenolypyruvate caboxykinase (PCK-2_Cys305) were all more reduced following exercise.

There were also 3 proteins containing Cys residues that become more oxidised in all 3 strains following exercise, Calreticulin (CRT-1_Cys133 (N2 and *skn-1*) and 101 (*prdx-2*), the basement membrane protein Papilin that contains a number of Cys residues oxidised after exercise (MIG-6 Cy1375 and 1523 (N2) Cys1161, 1408 and 1456 (*prdx-2*), Cys1375, 1408 and 1456 (*skn-1*) and Vitellogenin-2 (VIT-2_Cys228 and Cys1571). In both the N2 and *prdx-2* mutant strains following exercise that was 1 protein with a Cys residue more oxidised, Cytochrome oxidase assembly protein (COX-6b_Cys66). While in the *skn-1* and *prdx-2* strains, 5 proteins with specific Cys residues were oxidised in response to exercise, heat shock protein (HSP1_Cys243), actin-3 (ACT-3_Cys258), Cystatin (CPI-1_Cys99), Y37D8A.19_Cys54 and ZC373.2_Cys21/25.

It is notable that only in the N2 strain a number of Cys residues become more reduced which was not observed in either the *prdx-2* or *skn-1* mutants following exercise. Equally comparing the non-exercised strains N2 against the non-exercised *prdx-2* and *skn-1* mutant strains indicates that the Cys residues are basally more reduced in the *prdx-2* and *skn-1* mutants. The redox proteomics results would suggest that under basal conditions PRDX-2 is required for redox signalling and transfer of oxidative equivalents to target proteins, and PRDX-2 is essential for the signalling response to endogenous redox stress and downstream activation of SKN-1. The very distinct shift to a more oxidised profile would support the role of PRDX-2 as a key regulator of the intracellular redox environment with subsequent effects on endogenous redox signalling.

### Exercise improves locomotory activity in N2 strains but not prdx-2 or skn-1 mutants

In order to determine the physiological functional effects of exercise in wild type strains and mutant strains with an altered redox proteome, we used CeLeST (*C. elegans* swim tracking software) [43], to track the activity of the different strains following the 5day exercise protocol. N2 worms had increased wave initiation rate, activity index, travel speed and brush stroke compared to non-exercised controls (Fig.8). The *prdx-2* and *skn-1* mutants did not show a significant change in any of the 8 physiological parameters measured by the software in response to exercise compared to non-exercised controls. These results demonstrate the functional consequences of disrupted redox signalling as a result of loss of *prdx-2* and *skn-1* following an exercise protocol that can improve overall fitness and healthspan in wild type worms.

**Figure 8.**
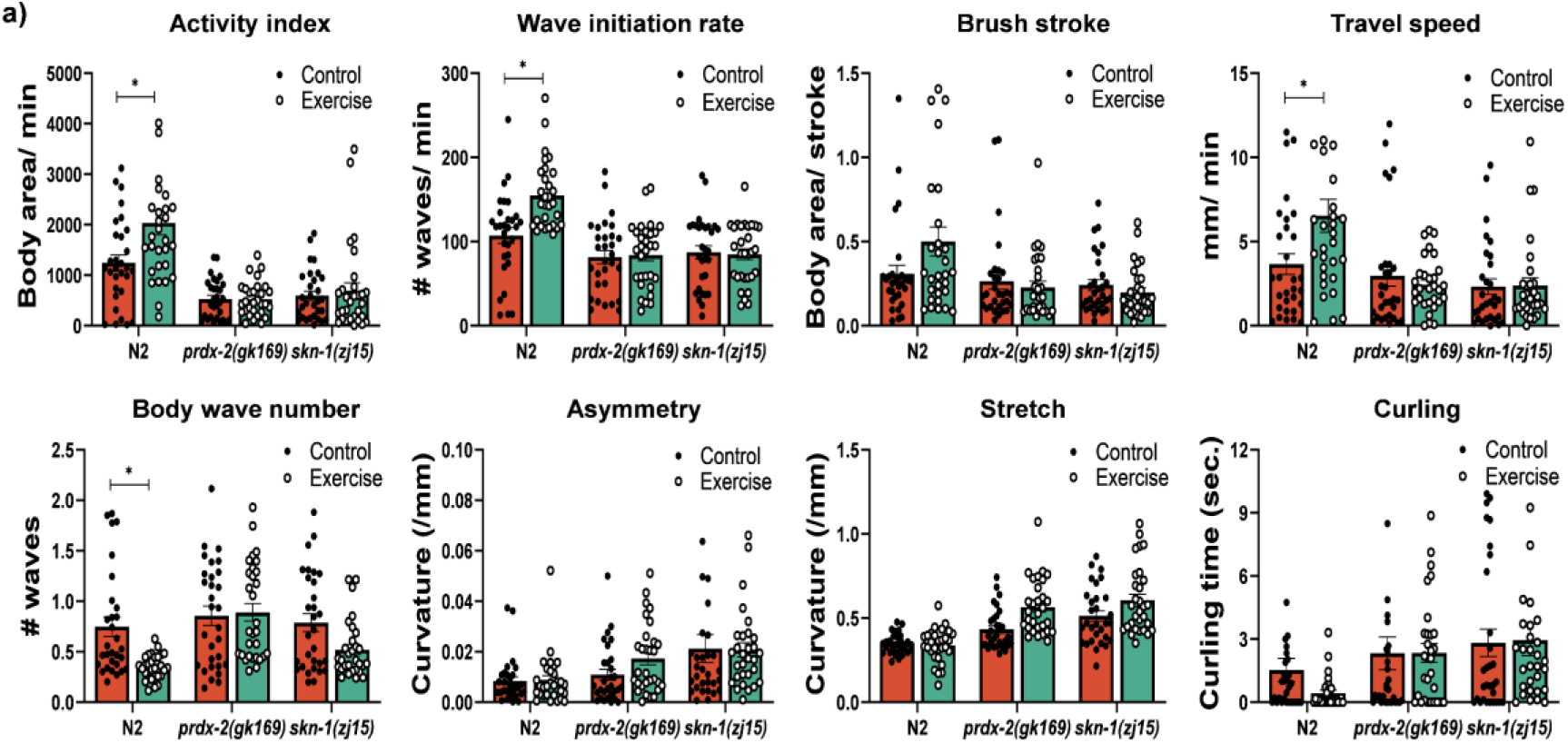
PRDX-2 and SKN-1 are required for improved fitness of worms following exercise. CeLeST analysis of physiological activity, activity index, wave initiation rate, travel speed, brush stroke, body wave number, asymmetry, stretch and curling in exercised and non-exercised worms. Experiments were performed with at least 30 worms per strain/experiment. *p values (Activity index: N2: Control vs Exercise= 0*.*0067; Wave initiation rate: N2: Control vs Exercise= 0*.*0002; Brush stroke: N2: Control vs Exercise= 0*.*0590; Travel speed: N2: Control vs Exercise= 0*.*0154; Body wave number: N2: Control vs Exercise= 0*.*0001; Curling: N2: Control vs Exercise= 0*.*0610)*.

In summary, our data demonstrate that a short physiological bolus addition of H_2_O_2_ result in a hormesis response in myoblasts through increasing antioxidant capacity and overall mitochondrial function, ultimately improving myogenesis. Transient knockdown of Prdx1 and Prdx2 suppressed this beneficial hormesis effect and led to smaller and fewer myotubes. Using *C. elegans*, a 5day swimming protocol in N2 wild type worms improved mitochondrial function and turnover, worms had increased ability to withstand external stress and prolonged lifespan. The swimming protocol did not have these beneficial effects in *prdx-2* mutants, there was no change in mitochondrial capacity, increased mitochondrial ROS, decreased ability to survive external stress and decreased lifespan following exercise. Similarly, there were no improvement in lifespan or survival following exercise in the *skn-1* mutants. The improved physiological fitness and activity following the exercise protocol in the N2 strain was not detected in the *prdx-2* or *skn-1* mutants. Furthermore, a redox proteomic approach demonstrated a more pronounced oxidation of specific Cys residues following exercise in the *prdx-2* and *skn-1* mutants. Together our data support the essential role of PRDX-2 in regulating redox signalling cascades following physiological endogenous ROS generation via exercise.

## Discussion

Exercise results in widespread changes in gene transcription and translation, however exactly how the mechanical events initiated during contractile activity are interpreted intracellularly leading to functional adaptations are unknown. Contraction of skeletal muscle generates endogenous ROS such as H_2_O_2_, required for adaptation to exercise activating precise signalling pathways, however ROS are relatively non-specific oxidising reagents [7, 34]. The abundance and kinetic reactivity of Prdxs to H_2_O_2_ suggest they can outcompete alternative redox signalling reactions [14]. A redox relay mechanism whereby the oxidative equivalents from H_2_O_2_ are transferred from evolutionary conserved Prxs to target proteins, would confer specificity on redox signalling. In this study, we have demonstrated that a short physiological bolus of H_2_O_2_ modifies the redox state of 2-Cys cytoplasmic Prdx1 and Prdx2. This is followed by an adaptive hormesis response in myoblasts, with increased mitochondrial capacity and improved myogenesis. Knockdown of 2-Cys cytoplasmic Prdxs prevents this adaptive response and although not essential for myogenesis under normal conditions, knockdown of Prdxs prior to H_2_O_2_ treatment results in dysregulated myogenesis as a result of disrupted redox signalling. Here, we implemented a 5day exercise protocol in *C. elegans* that promotes mitochondrial capacity, results in increased ability to withstand external stressors, improved overall fitness and extends lifespan. In worms with non-functional PRDX-2, these beneficial adaptive responses to exercise were not apparent. In contrast, the *prdx-2* mutant strain had decreased ability to survive external stress and shortened lifespan following the exercise protocol, indicating PRDX-2 is essential for adaptive redox signalling in response to endogenous stress. Prdxs role in the adaptive response to a disrupted redox environment is via reversible oxidation of the peroxidatic Cys residue and has been suggested to transfer oxidising equivalents to target proteins [12, 16, 18]. Our redox proteomic approach identified a shift in the redox state of individual Cys residues to a relatively more oxidised redox state of Cys residues following an exercise intervention in *prdx-2* mutants, supporting the role of PRDX-2 in the regulation of the intracellular redox environment following endogenous ROS generation.

The hormesis effect of ROS has been widely reported (for review see [44, 45], low level exposure of cells to ROS can induce beneficial adaptive responses including improved antioxidant defence mechanisms and mitochondrial biogenesis, while high levels can be detrimental inducing oxidative damage. The response of myoblasts to a short bolus addition of H_2_O_2_ appears to prime the cells for improved myogenesis, associated with increased nuclear localisation of key antioxidant transcription factors, increased expression of antioxidant proteins and proteins involved in mitochondrial turnover, along with improved mitochondrial capacity. Our data indicate that the 2-Cys Prdxs are essential for the hormesis response; this beneficial adaptive response was suppressed in myoblasts with decreased expression of Prdx1 and Prdx2, resulting in disrupted myogeneis. Similarly in *C. elegans* the failure of the *prdx-2* mutant strain to demonstrate the beneficial adaptation to the exercise indicates that PRDX-2 is essential for hormesis and endogenous redox signalling. Previously, the role of PRDX-2 has been reported to be required for the lifespan extension of *C. elegans* cultivated at lower temperatures and in response to metformin treatment [29, 30], interventions that affect the endogenous redox environment and activation of an adaptive redox response. PRDX-2 provides an essential link between the activation of SKN-1 via p38 MAPK and insulin signalling [31, 46].

The global proteomic approach revealed some very distinct changes in the proteomes following exercise, N2 strains had increased expression of proteins that are typically detected in mammalian studies of exercise with increased expression of key metabolic proteins and those involved in fatty acid beta oxidation. The proteomic changes in response to exercise are very distinct between the different strains, with a significant increase in abundance of ion transporters and downregulation of cuticle proteins in the wild type N2 strain. Interestingly, the *prdx-2* mutants in response to exercise increased the abundance of many cytoskeletal related proteins and also GSK-3, an inhibitor of SKN-1 activation [47]. Also of interest was a decrease in abundance of proteins related to miRNA biogenesis and proteins involved in the transsulfuration pathways. In the *skn-1* mutant there was a notable increase in the abundance of a number of iron regulatory proteins following exercise. The global proteomic analysis also revealed some common changes in all strains following exercise, AKT-1 and DUF148 increased in abundance. AKT-1 is involved in the phosphorylation of DAF-16 and SKN-1, inhibiting their translocation to the nucleus [39, 48]. Similarly, HACD-1 and PMT-1 decreased in abundance following exercise in the 3 strains. HACD-1 is predicted to be located in mitochondria and involved in fatty acid oxidation while PMT-1 is involved in phosphotidylcholine synthesis, a critical component of the plasma membrane [49]. These conserved changes across strains would suggest a non-redox dependent metabolic response to exercise.

Redox signalling via thiol disulphide exchange on Cys residues has been reported to regulate a host of signalling pathways for physiological adaptations and pathologies. Cys residues have the most extreme conservation pattern within proteins, highly conserved when they form part of an active site or involved in co-factor binding and poorly conserved otherwise [50]. Our redox proteomic approach identified a large number of redox sensitive Cys residues that have previously been reported in *C. elegans* to respond to changes to external H_2_O_2_ [51] and in insulin signalling mutants [52]. We detected Cys366 from mitochondrial complex I subunit NDUS2 as redox sensitive, although it did not significantly change in response to exercise, reversible oxidation of this residue has been reported to determine the behavioural response to hypoxia [53]. Subtle changes in the reversible oxidation of specific Cys residues have been reported to induce widespread physiological effects, for example oxidation of Akt1 Cys60 from 0.5% to 1.25% following insulin stimulation is required for phosphatidylinositol (3,4,5)-trisphosphate dependent recruitment of Akt1 to the plasma membrane [54]. The redox sensitive proteins identified here correspond to a large number of human orthologue proteins previously identified as interacting with human 2-Cys Prdx’s [55]. We also demonstrate that mutants lacking functional PRDX-2 were characterised by a more reduced redox state of Cys residues compared to N2 in non-exercised strains but that in response to exercise the *prdx-2* mutant had a more oxidised redox profile of Cys residues. Similarly, there was an increase in mitochondrial ROS following exercise in *prdx-2* strains accompanied by decreased lifespan and survival to external stressors.

Prdx2 has previously been identified as interacting with STAT3, forming an inter-disulphide bond following a brief exposure to H_2_O_2_, attenuating its activity in HeLa cells [12] but the precise mechanism of Prdx2 role in a redox signalling cascade requires further study. Membrane molecular chaperones may facilitate the redox relay mechanism of target proteins interacting with Prdxs close to the endogenous source of ROS. A recent study indicates Prdx2 forms a scaffold with Annexin A2 via a disulphide bridge facilitating STAT3 interactions [56]. In muscle, Annexin 2A is required for the acute inflammatory response to facilitate plasma membrane repair in injured muscle following disruption of the mitochondrial architecture, ROS and calcium release [57], which would support its role in a redox-dependent signalling pathway. We did not identify significant changes in the redox state of *C. elegans* orthologues of Annexins (NEX-1_Cys112 and NEX-3_Cys196) but did detect significant changes in the redox state of hsp chaperone protein (HSP-1). Other proteins of particular interest with Cys specific redox changes include calcium handling proteins CPN-4, CRT-1 and EAT-6 that might coincide with the changes in overall abundance of proteins involved in ion transport in the N2 strain following exercise but not detected in the *prdx-2* mutant strain. A number of specific Cys residues become relatively more reduced following exercise only in the N2 strain, particularly proteins involved in mitochondrial metabolic pathways. This would support the role of PRDX-2 as required for effective redox signalling following endogenous ROS generation during exercise and the adaptive response to exercise.

The redox state of Prdxs has been linked to cellular metabolism in a variety of different cell models, the circadian cyclical oxidation of Prdxs in red blood cells provides a link between the central core clock and peripheral clocks [58]. In yeast, the oxidation state of cytoplasmic 2-Cys Prdxs regulates the ultraradian cycles between high and low oxygen consumption and ultimately determines timely cell division [18]. Exercise induced endogenous generation of ROS could potentially act as a Zeitgeber in localised ROS generation via the reversible oxidation of cytoplasmic Prdxs with subsequent effects on transcription factor activation, gene expression and the adaptive response to exercise. Interestingly, worms that have an acute early life increase in ROS have improved epigenetic responses, stress resistance and longevity as a result of mitohormesis [59]. This would suggest that it is possible to individualise longevity by manipulation of the redox environment via acute endogenous ROS generation such as in exercise as demonstrated here or in situations with disrupted redox signalling, that result in altered mitochondrial dynamics such as ageing or chronic disease.

The body wall muscle of the nematode *C. elegans* closely resembles the sarcomere of striated muscle in vertebrates [60] and with age there is disrupted mitochondrial dynamics, lipid accumulation and loss of muscle mass [26, 32]. *C. elegans* also contains orthologues of conserved redox sensitive proteins described in the mammalian response to exercise. Additionally, the improvements in mitochondrial capacity, fitness and longevity are consistent in the wild type strains compared to mammalian exercise studies. Due to technical limitations, it was not possible to perform the proteomic approach from isolated body wall muscle of *C. elegans*. However, the expression of PRDX-2 is relatively high in all tissues, particularly in neurons and the redox proteomic approach would support the key signalling effects of PRDX-2 in the whole organism. The major advantage of the redox proteomic approach is that both overall protein abundance and relative quantification of individual Cys residues can be obtained in a single analysis, alternative approaches that use affinity-based purification of modified Cys residues can have increased sensitivity but lose information on overall protein abundance.

## Conclusion

Our data support the role of 2-Cys Prdxs as required for hormesis in response to endogenous ROS generation such as during exercise. A specific exercise regime results in redox adaptive responses as a result of acute ROS generation and the associated signalling events resulting in improved mitochondrial function that will increased healthspan in wild type worms. Worms deficient for PRDX-2 did not achieve the beneficial adaptive response to exercise, in contrast exercise had a detrimental effects on the *prdx-2* strain. Underlying this response there was a shift to a relatively more oxidised state in the *prdx-2* mutant strain as indicated by the redox proteomic approach, while in wild type worms specific Cys residues were both more reduced and oxidised. Together this data demonstrates the key role of PRDX-2 in the regulation of the intracellular redox environment required for the adaptive hormesis response to endogenous ROS generation during exercise.

### Reagents and Resources

**Reagents and Resources:**
| REAGENT or RESOURCE | SOURCE | IDENTIFIER |
| --- | --- | --- |
| **Antibodies** |  |  |
| rabbit anti-Peroxiredoxin 1 | Abcam | Cat# ab109498,<br>RRID:AB_10865914 |
| rabbit anti-Peroxiredoxin 2 | Cell Signaling Technology | Cat# 46855,<br>RRID:AB_2799310 |
| rabbit anti-Peroxiredoxin 2 (C. elegans) | Elizabeth Veal lab [28] | N/A |
| mouse anti-Peroxiredoxin 3 | Abcam | Cat# ab16751,<br>RRID:AB_302498 |
| mouse anti-Peroxiredoxin 5 | Abcam | Cat# ab16944,<br>RRID:AB_2168508 |
| rabbit anti-Peroxiredoxin 6 | Abcam | Cat# ab133348,<br>RRID:AB_11155931 |
| rabbit anti-Peroxiredoxin-SO3 | Abcam | Cat# ab16830,<br>RRID:AB_443491 |
| rabbit anti-Nrf2 | Abcam | Cat# ab31163,<br>RRID:AB_881705 |
| rabbit anti-NF-κB p65 | Abcam | Cat# ab16502,<br>RRID:AB_443394 |
| rabbit anti-FoxO3a | Cell Signaling Technology | Cat# 2497, RRID:AB_836876 |
| mouse anti-STAT3 | Cell Signaling Technology | Cat# 9139, RRID:AB_331757 |
| mouse anti-Phospho-Stat3 | Cell Signaling Technology | Cat# 9138, RRID:AB_331262 |
| rabbit anti-SOD2 | Abcam | Cat# 2299-1,<br>RRID:AB_1267348 |
| rabbit anti-Thioredoxin 1 | Abcam | Cat# ab86255,<br>RRID:AB_1925501 |
| mouse anti-Thioredoxin 2 | Abcam | Cat# ab16857,<br>RRID:AB_302548 |
| rabbit anti-TOM20 | Cell Signaling Technology | Cat# 42406,<br>RRID:AB_2687663 |
| rabbit anti-PGC-1α | Cell Signaling Technology | Cat# 2178, RRID:AB_823600 |
| rabbit anti-TFAM | Abcam | ab272885 |
| rabbit anti-ULK1 | Abcam | Cat# ab128859,<br>RRID:AB_11156928 |
| rabbit anti-LC3b | Abcam | Cat# ab192890,<br>RRID:AB_2827794 |
| rabbit anti-SQSTM1/p62 | Cell Signaling Technology | Cat# 88588,<br>RRID:AB_2800125 |
| mouse anti-BNIP3 | Abcam | Cat# ab10433,<br>RRID:AB_2066656 |
| rabbit anti-PINK1 | Abcam | Cat# ab23707,<br>RRID:AB_447627 |
| rabbit anti-Parkin | Abcam | Cat# ab15954,<br>RRID:AB_443270 |
| rabbit anti-PARK7/DJ1 | Abcam | Cat# ab76241,<br>RRID:AB_2283514 |
| Goat anti-Rabbit IgG(H+L) Alexa Fluor™ 546 | Thermo Fisher Scientific | Cat# A-11035,<br>RRID:AB_2534093 |
| Goat anti-Mouse IgG(H+L) Alexa Fluor™ Plus 555 | Thermo Fisher Scientific | Cat# A32727,<br>RRID:AB_2633276 |
| mouse anti-MF20 | Developmental Studies Hybridoma Bank | Cat# MF 20, RRID:AB_2147781 |
| rabbit anti-Keap1 | Abcam | Cat# ab66620, RRID:AB_1141055 |
| IRDye 800CW Goat anti-Rabbit IgG | LI-COR Biosciences | Cat# 926-32211, RRID:AB_621843 |
| IRDye 800CW Goat anti-Mouse IgG | LI-COR Biosciences | Cat# 925-32210, RRID:AB_2687825 |
| **Bacterial and Virus Strains** |  |  |
| *E. coli*: Strain OP50 | Caenorhabditis Genetics Center | N/A |
| **Chemicals, Peptides, and Recombinant Proteins** |  |  |
| DMEM | Sigma | Cat# D5796 |
| PBS | Sigma | Cat# D8537 |
| FBS | Sigma | Cat# F7524 |
| HS | Thermo Fisher Scientific | Cat# 26050088 |
| Penicillin/Streptomycin | Thermo Fisher Scientific | Cat# 15140122 |
| TrypLE™ Express Enzyme | Thermo Fisher Scientific | Cat# 12604013 |
| H<sub>2</sub>O<sub>2</sub> | Sigma | Cat# H1009 |
| NaCl | Sigma | Cat# S9888 |
| Trizma® base | Sigma | Cat# T1503 |
| EDTA | Sigma | Cat# ED2SS |
| SDS | Sigma | Cat# L3771 |
| Glycine | Sigma | Cat# G8898 |
| Triton™ X-100 | Sigma | Cat# T8787 |
| NEM | Sigma | Cat# E3876 |
| Beta-mercaptoethanol | Sigma | Cat# M7522 |
| Bradford Reagent | Bio-Rad | Cat# 5000006 |
| BSA | Sigma | Cat# A3059 |
| Glycerol | Sigma | Cat# G6279 |
| Bromophenol blue | Sigma | Cat# 114391 |
| Acrylamide | Sigma | Cat# A3699 |
| APS | Sigma | Cat# A3678 |
| TEMED | Sigma | Cat# T9281 |
| Methanol | Sigma | Cat# 34860 |
| Ethanol | Sigma | Cat# E7023 |
| Lipofectamine 2000 Reagent | Thermo Fisher Scientific | Cat# 11668019 |
| Nystatin | Sigma | Cat# N3503 |
| Paraquat | Sigma | Cat# 856177 |
| TBHP | Sigma | Cat# 654833 |
| DAPI | Sigma | Cat# D9542 |
| TRIzol Reagent | Life Technologies | Cat# 15596018 |
| Chloroform | Sigma | Cat# C0549 |
| Isopropanol | Sigma | Cat# I9516 |
| Protease Inhibitor Cocktail | Sigma | Cat# P8340 |
| SYBR Green | Qiagen | Cat# 339347 |
| RNase free water | Sigma | Cat# W4502 |
| Hydromount | Scientific laboratory supplies | Cat# D2176 |
| Urea | Sigma | Cat# U5378 |
| Agar | Sigma | Cat# A1296 |
| Peptone | Sigma | Cat# 91249 |
| Cholesterol | Sigma | Cat# C8667 |
| CaCl<sub>2</sub> | Sigma | Cat# C1016 |
| MgSO<sub>4</sub> | Sigma | Cat# M7506 |
| KH<sub>2</sub>PO<sub>4</sub> | Sigma | Cat# P9791 |
| K<sub>2</sub>HPO<sub>4</sub> | Sigma | Cat# 3786 |
| Na<sub>2</sub>HPO<sub>4</sub> | Sigma | Cat# S0876 |
| NaOH | Sigma | Cat# S5881 |
| NaClO | Sigma | Cat# 1056142500 |
| Nitrocellulose Blotting membrane | GE Healthcare Life Sciences | Cat# 1060003 |
| Ponceau S Staining Solution | Thermo Fisher Scientific | Cat# A40000279 |
| Acetic acid | Sigma | Cat# A6283 |
| **Critical Commercial Assays** |  |  |
| MitoTracker™ Red CMXRos | Thermo Fisher Scientific | Cat# M7512 |
| MitoTracker™ Green FM | Thermo Fisher Scientific | Cat# M7514 |
| MitoSOX™ Red mitochondrial superoxide indicator | Thermo Fisher Scientific | Cat# M36008 |
| Cell Mito Stress Test Kit | Agilent Technologies | Cat# 103015-100 |
| **Deposited Data** |  |  |
| PRIDE | ProteomeXchange Consortium | PXD037698 |
| **Experimental Models: Cell Lines** |  |  |
| C2C12 | ATCC | Cat # CRL-1772 |
| **Experimental Models: Organisms/Strains** |  |  |
| *C. elegans*: wild type | Caenorhabditis Genetics Center | N2 |
| *C. elegans*: *prdx-2(gk169)* II | Elizabeth A Veal and Caenorhabditis Genetics Center | VC289 |
| *C. elegans*: *skn-1(zj15)* IV | Caenorhabditis Genetics Center | QV225 |
| *C. elegans*: *bli-3(e767) lin-17(n677)* I | Caenorhabditis Genetics Center | MT1344 |
| *C. elegans*: *dvIs19[(pAF15)gst-4p::gfp]* III | Caenorhabditis Genetics Center | CL2166 |
| *C. elegans*: *zcls14 [myo-3::gfp(mit)]* | Caenorhabditis Genetics Center | SJ4103 |
| *C. elegans*: *I (unc-119(ed3); Ex[myo-3p tomm20::Rosella;unc-119(+)]* | Tavernarakis Lab | IR2539 |
| **Oligonucleotides** |  |  |
| mouse Prdx1 siRNA | Life Technologies | Cat# 4390824 |
| mouse Prdx2 siRNA | Life Technologies | Cat# 4390824 |
| **Software and Algorithms** |  |  |
| Prism 7 | GraphPad Software | RRID:SCR_002798 |
| ImageJ | NIH | RRID:SCR_003070 |
| Seahorse Wave | Agilent Technologies | RRID:SCR_014526 |
| Image Studio Lite | Image Studio Lite | RRID:SCR_013715 |
| CeLeSt | Christophe Restif et al.,<br>2014 [43] | N/A |

## Methods

### Cell culture

C2C12 skeletal muscle myoblasts were maintained in GM (growth medium) containing DMEM (Dulbecco’s Modified Eagle’s Medium, high glucose), 10% FBS (Fetal bovine serum) and 1% P/S (Penicillin/Streptomycin) at 37°C in 5% CO_2_ [61]. For H_2_O_2_ treatments, 70-80% confluent C2C12 myoblasts were treated with a range of H_2_O_2_ concentrations for 10 mins and a time course of 25 μM H_2_O_2_. C2C12 myoblasts were immediately homogenised in an alkylating lysis buffer (150mM NaCl, 20mM Tris pH 7.5, 1mM EDTA pH 8.3, 0.5% SDS, 1% Triton, 100mM NEM) to prevent thiol disulphide exchange [42]. For the adaptive response to physiological H_2_O_2_, 50-60% confluent C2C12 myoblasts were treated with 25 μM H_2_O_2_ for 10 mins, the media was exchanged for fresh media and cells were allowed to grow for 3h-24h, proteins were isolated with alkylating lysis buffer containing protease inhibitor cocktail or immunostained. For analysis of myotube formation, C2C12 myoblasts following 10 mins of 25 μM H_2_O_2_ treatment and 24h recovery in GM, cells were maintained in DM (differentiation medium) containing DMEM, 2% HS (horse serum) and 1% P/S. Protein was isolated during the specified times during differentiation process, immunostaining of myotube was performed 5 days after initiation of differentiation. For the transient knockdown of Prdx1 and Prdx2, 20-30% confluent C2C12 myoblasts were treated with 50nM of siPrdx1 and siPrdx2 for 5-6 h, medium was refreshed with GM for 2 days previous to treatment with 10mins of 25μM H_2_O_2_, proteins were isolated 24h after treatment or immunostained as described below.

### *C. elegans* strains

*C. elegans* were cultured on NGM plates seeded with *E. Coli* (OP50) at 20 °C. C. elegans stains N2 wild type, QV225 (*skn-1(zj15)* IV), MT1344 (*bli-3(e767) lin-17(n677)* I), CL2166 (dvIs19[*(pAF15)gst-4p::gfp*] III) and SJ4103 (zcIs14 [*myo-3::gfp(mit)*]) strains were obtained from the Caenorhabditis Genetics Center (CGC) funded by NIH Office of Research Infrastructure Programs (P40 OD010440). The VC289 (*prdx-2(gk169)* II) strain was a gift of Elizabeth A Veal (Newcastle University, UK). The IR2539 (*unc-119(ed3); Ex*[*pmyo-3 tomm-20::Rosella;unc-119*(+)] was a gift from the Tavernarakis lab University of Crete, Greece.

### Western blotting

Protein extracts from cells and *C. elegans* were calculated using Bradford reagent following homogenisation in an alkylating lysis buffer. For Western blotting, 20 μg of protein was loaded on 12% reducing or non-reducing SDS PAGE gels. Proteins were transferred using a semi-dry blotter and membrane was stained with Ponceau S for normalisation. Following washing membranes were blocked in 5% milk in TBS-T for 1 h at room temperature and membranes were incubated with primary antibodies (see material tables) with a dilution of 1:1000 in 5% milk for overnight. Membranes were washed 3 × 10 min in TBS-T and incubated with secondary antibody with a dilution of 1:10000 in TBS-T in the dark for 1h and images were acquired using Odyssey Fc imaging system. Quantification and normalization of blots were analysed using Image Studio Lite.

### Cell Microscopy

Immunocytochemistry of C2C12 myoblasts, cells were cultured with laminin covered coverslips in 10% GM until 50-60% confluence. Following 10 min of 25 μM H_2_O_2_ treatment, media was changed and cells were allowed to grow for 3h-24h. Samples were fixed with ice cold methanol for 5 min and then in blocking buffer (1% BSA, 10 % Horse serum, 0.3M Glycine, 0.2% Triton X in PBS) for 1 h RT. Cells were incubated with primary antibodies (NRF2, NF-κB, FOXO3a, STAT3) with a dilution of 1:1000 in 5% HS for overnight. Next, cells were washed 3 × 5 min in PBS and incubated with secondary antibody (1:2000 dilution in PBS) in the dark for 1h followed by incubated with DAPI for 10 min. One drop of hydromount was used and left overnight at 4°C and images were acquired using EVOS M7000. Quantification of images were performed using ImageJ [62].

For the MF 20 immunostaining of myotubes, 24h after 10 min of 25 μM H_2_O_2_ treatment, media was changed with DM to allow the differentiation to myotubes, MF20 immunostaining was performed as described after 5 days, images were acquired using EVOS M7000 and quantification of images were performed using ImageJ [63].

For MitoTracker and MitoSOX staining of myoblasts, 24h after the H_2_O_2_ treatment, medium was removed and washed with PBS, myoblasts were incubated with Hoechst 33342 for 5 min, washed with PBS and incubated with 200 nM MitoTracker Green solution for 30 min, washed with PBS and incubated with 5 μM MitoSOX Red solution for 10 min, washed with PBS and images were taken by EVOS M7000 at 60× magnification. The fluorescence intensity per each cell was assessed by ImageJ [64].

### Swimming exercise

Swimming exercise protocol was performed according to [27]. Briefly, synchronized worms were bleached by sodium hydroxide bleaching buffer, followed by overnight incubation in M9 buffer on a rocker at 20 °C. The populations of L1 worms were then transferred to 60 mm NGM plates seeded with OP50 for 2 days to obtain adult D1 worms. Worms were washed off plates with 3 mL M9 buffer and allowed to settle under gravity, the supernatant including OP50 and larvae removed and repeated 3 times. Worms were transferred to unseeded NGM plates with M9 buffer (exercise group) or unseeded NGM plates (control group) using a glass Pasteur pipette and all plates moved to 20 °C incubator for 90 min. After swimming exercise, worms from both conditions were washed off with M9 buffer, gravity settled and transferred to new 60 mm NGM plates seeded with OP 50 in 20 °C incubator. Swimming exercise were performed 2×90 min at 9:00 AM and 3:00 PM daily for 5 days.

### Lifespan assay

Lifespan assay was performed after the last day of swimming exercise, 105 worms were separately transferred to 3 seeded NGM plates. The worms were transferred lightly by a picker to new plates every 1-2 days until no larvae hatched. Death of worms was considered when no response by a light tap with a picker, survival was assessed every day.

### Oxidative stress assays

One day after the swimming exercise regimen, the paraquat stress assay was performed according to [65]. A fresh vial of 100 mM Paraquat was prepared in M9 buffer and pipetted 60 μL into wells of a 96-well plate. 10 worms per each well, 80 worms were picked and transferred into 8 wells per each condition. The TBHP assay was performed according to [66]. TBHP plates were prepared one day before experiment and placed in an airtight box. 25 worms per plate, 75 worms were picked onto TBHP plates per each condition. Death of worms was considered when no response by a gentle tap with a picker, survival was assessed every 1-2 h.

### Microscopy of C. elegans

For SJ4103 (zcIs14 [*myo-3::gfp(mit)*]) strain, the imaging was performed one day after the swimming exercise regimen using EVOS M7000 at 60× magnification according to [41]. 45 worms per each condition were immobilized by 20 mM Levamisole on the agar slide and covered with coverslip. The mitochondrial morphology between pharynx and vulva was assessed based on three categories: filamentous (filamentous mitochondrial network), punctate (most fragmented mitochondria), intermediate (isolated mitochondrial network).

Mitochondrial dynamics was assessed using (*unc-119(ed3);* Ex[*pmyo-3 tomm-20::Rosella;unc-119*(+)] immobilization of worms was performed described above and 60 worms were imaged using EVOS M7000 at 60× magnification, the ratio of green versus red fluorescence intensity in a representative head region per each worm was assessed by ImageJ [40]. SKN-1 Transcriptional activated was assessed using CL2166 (dvIs19[(*pAF15)gst-4p::gfp* III) strain, 5 worms per group and 45 worms in total imaged for each condition, worms were immobilized on unseeded NGM plates and imaged by EVOS at 10× magnification. The GFP fluorescence intensity per each worm was assessed by ImageJ.

For the MitoTracker and MitoSOX immunostaining of *C. elegans*, 45 worms were imaged for each condition, worms were incubated with 50 μL of 2.5 μM MitoTracker Red CMXRos or 10 μM MitoSOX Red diluted in M9 buffer for 10 min or 1 h at 20°C in the dark. The worms were transferred to seeded NGM plates for 2 h at 20°C in the dark to avoid the accumulation of the staining in the guts of the nematode, then 5 worms per group were immobilized on the unseeded NGM plates and imaged by EVOS at 10× magnification. The fluorescence intensity per each worm was assessed by ImageJ [64].

### Mitochondrial respiration rates

C2C12 myoblast were performed in triplicate and seeded in XF HS Mini Analyzer plates at a density of 8000 cells per well, total volume 200 μL/ well, and incubated at 37°C in 5% CO2 for 24 h. Cells were treated with 25 μM H_2_O_2_ for 10 min and 10% GM was refreshed to recover for 24 h. Cells were washed once with XF Real-Time Mito-Stress Assay Medium (10 mM glucose, 1 mM sodium pyruvate, and 2 mM glutamine, pH 7.4), medium was refreshed and cells were incubated at 37°C in 0% CO2 for 1h. The compounds (2 μM Oligomycin, 2 μM FCCP and 1 μM antimycin A) were added in the hydrated cartridge, then the cartridge was loaded with the plate and calibration process in the XF HS Mini Analyzer was performed.

For mitochondrial respiration rates of *C. elegans*, the compounds (10 μM FCCP and 40 mM sodium azide) were added in the hydrated cartridge, 3 replicates and 8-13 worms per replicate were transferred into a XF HS Mini Analyzer plate before the calibration process started [67].

### CeleST

CeleST (*C. elegans* Swim Test) was conducted as described by [43]. One day after the swimming exercise regimen, a 10 mm pre-printed ring on the surface of a glass microscope slide was used and covered with 50 μL M9 Buffer. 4-5 worms per each group and 30 worm totally were picked and placed into the swimming area, 60-s movies with ∼16 frames per second of the animals were taken using a Nikon LV-TV microscope at 1× magnification with a OPTIKA C-P20CM camera.

### Mass Spectrometry

Protein extracts were diluted to a final concentration of 2M Urea 100 mM Tris-HCl pH 8. In-solution digestion, using Lys-C (Wako) and trypsin (Trypzean, Sigma) was performed during 10h at 37°C (protein:enzyme ratio 1:50). Resulting peptides were desalted using a C18 plate.

LC-MS/MS was done by coupling an UltiMate 3000 *RSLCnano* LC system to a Orbitrap Exploris™ 480 mass spectrometer (Thermo Fisher Scientific). Peptides were loaded into a trap column (Acclaim™ PepMap™ 100 C18 LC Columns 5 μm, 20 mm length) for 3 min at a flow rate of 10 μl/min in 0.1% FA. Then, peptides were transferred to an EASY-Spray PepMap RSLC C18 column (Thermo) (2 μm, 75 μm x 50 cm) operated at 45 °C and separated using a 60 min effective gradient (buffer A: 0.1% FA; buffer B: 100% ACN, 0.1% FA) at a flow rate of 250 nL/min. The gradient used was, from 2% to 6% of buffer B in 2 min, from 6% to 33% B in 58 minutes, from 33% to 45% in 2 minutes, plus 10 additional minutes at 98% B. Peptides were sprayed at 1.5 kV into the mass spectrometer via the EASY-Spray source and the capillary temperature was set to 300 °C.

The mass spectrometer was operated in a data-dependent mode, with an automatic switch between MS and MS/MS scans using a top 18 method. (Intensity threshold ≥ 3.6e5, dynamic exclusion of 20 sec and excluding charges +1 and > +6). MS spectra were acquired from 350 to 1400 m/z with a resolution of 60,000 FWHM (200 m/z). Ion peptides were isolated using a 1.0 Th window and fragmented using higher-energy collisional dissociation (HCD) with a normalized collision energy of 29. MS/MS spectra resolution was set to 15,000 (200 m/z). The normalised AGC ion target values were 300% for MS (maximum IT of 25 ms) and 100% for MS/MS (maximum IT of 22 msec).

Raw files were processed with MaxQuant (v 1.6.0.16) using the standard settings against a *C. elegans* protein database (UniProtKB/Swiss-Prot/TrEMBL, 27385 sequences). N-ethylmaleimide and D5 N-ethylmaleimide on cysteines, oxidation of methionines and protein N-term acetylation were set as variable modifications. Minimal peptide length was set to 7 amino acids and a maximum of two tryptic missed-cleavages were allowed. Results were filtered at 0.01 FDR (peptide and protein level).

Afterwards, the “proteinGroups.txt” file was loaded in Prostar [68] using the intensity values for further statistical analysis. Briefly, proteins with less than three valid values in at least one experimental condition were filtered out. Then, a global normalization of log2-transformed intensities across samples was performed using the LOESS function. Missing values were imputed using the algorithms SLSA for partially observed values and DetQuantile for values missing on an entire condition. Differential analysis was done using the empirical Bayes statistics Limma. Proteins with a p value < 0.01 and a Log_2_FC ratio >0.58 or <-0.58 were defined as significantly changed. The FDR was estimated to be below 6.5% by Benjamini-Hochberg. Analysis of the redox dependent changes was performed as described above only Cys containing residues labelled with light or heavy NEM were filtered from original data set. Cys residues with a p value < 0.05 and log2 ratio >1.5 or <-.1.5 were considered significantly changed in response to exercise. Proteomic data has been deposited in the PRIDE repository accession PXD037698.

### Statistical analysis

Images were quantified semi-automatically by ImageJ and Image Studio Lite followed by manual correction [69]. For Western blot images, the band intensities were normalized according to protein intensity from Ponceau S staining. For the microscopy of C2C12, at least 3-6 images were captured randomly at 10× or 40× magnification from different fields of view per each biological replicate. For the exercise related transcription factors analysis, percentage of nuclear localization among total cells were assessed. For myogenic differentiation analyses (MF20 immunostaining), in each field of view, average diameter of all myotubes was measured as myotube diameter, average area fraction of myotubes was calculated as myotube area, percentage of nuclear contained within myotubes to the total number of nuclear was assessed as fusion index. For the MitoTracker and MitoSOX immunostaining, the relative fluorescence intensity per each cell were assessed. For the microscopy of *C. elegans*, at least 45 worms were measured at 10× or 60× magnification from different fields of view per each condition. For SJ4103 strain, mitochondrial morphology between pharynx and vulva was assessed based on three categories: filamentous (filamentous mitochondrial network), punctate (most fragmented mitochondria), and intermediate (isolated mitochondrial network). For the mitochondrial dynamics the ratio of green versus red fluorescence was assessed [40]. For CL2166 strain, MitoTracker and MitoSOX immunostaining, fluorescence intensity per each worm was assessed. Details of the statistical analyses performed are indicated in the corresponding figure legend. Student t-test was used for the analysis of statistical differences between two groups. One-way or two-way ANOVA was performed to compare more than two groups. Log-rank test was used for the comparation of lifespan and tolerance of oxidative stress assay. Chi-square test was used to compare the distribution into multiple categories. p-value < 0.05 was considered statistically significant. Statistical analysis and graphs were performed using GraphPad Prism 7. All source data used to produce the figures in this manuscript are available in the source data file.

## Acknowledgements

We would like to sincerely thank Elizabeth Veal lab for comments on manuscript, providing the anti-PRDX-2 antibody and VC289 *prdx-2* mutant strain and the Tavernarakis lab for kindly providing the IR2539 mitophagy reporter strain. QX studentship is funded by the Chinese Scholarship Council (CSC), JCCM studentship is funded by the College of Nursing Medicine and Health Sciences, University of Galway. The mass spectrometry work was funded by European Proteomics Infrastructure Consortium providing access (EPIC-XS): Project number 823839.

## Author Contributions

Conceptualisation, BMcD; methodology QX, JCCM, KW, EZ, JM, AVM and BMcD; investigation, QX, JCCM, EZ and BMcD; resources, KW, JM, AVM and BMcD; writing-original draft BMcD; writing-review and editing, QX, JCM, EZ, JM, AVM, KW and BMcD; supervision, KW and BMcD.

**Supplementary Figure 1.**
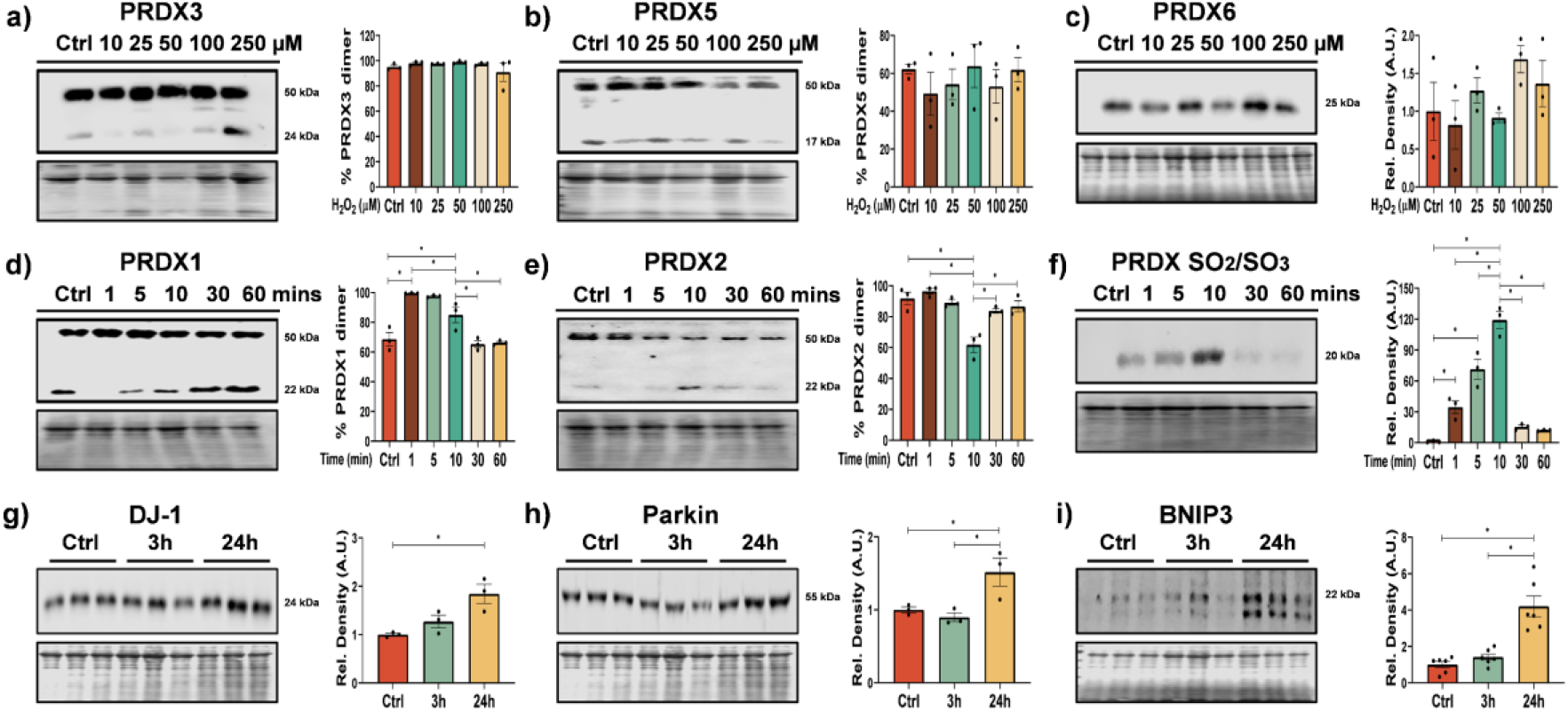
Effects of different concentrations of H_2_O_2_ on monomer/dimer ratio analysed by immunoblotting for Prdx3 (a), Prdx5 (b), and Prdx6 (c) in C2C12 myoblasts 25 μM H_2_O_2_ for 10 min. Immunoblotting for monomer/dimer of PRDX1 (d) and PRDX2 (e) and Prdx SO_2_/SO_3_ (f) following 25 μM H_2_O_2_ treatment for 1, 5, 10, 40 or 60 min. Immunoblotting of proteins involved in mitochondrial turnover (Protein DJ-1, Parkin and Bnip3) in C2C12 myoblasts treated with 25 μM H_2_O_2_ for 10 min and allowed to proliferate for 3h and 24h. Graphs are the mean +/-SEM and all experiments were performed with at least n=3-6, one-way ANOVA was used for significance between groups and *p-*value of <0.05 was considered as statistically significant *(*p*<0.05). *p values (d: Ctrl vs 10 mins= 0*.*0248; e: Ctrl vs 10 mins= 0*.*0003; f: Ctrl vs 10 mins< 0*.*0001; g: Ctrl vs 24h= 0*.*0117; h: Ctrl vs 24h= 0*.*0495; i: Ctrl vs 24h< 0*.*0001)*.

**Supplementary Figure 2.**
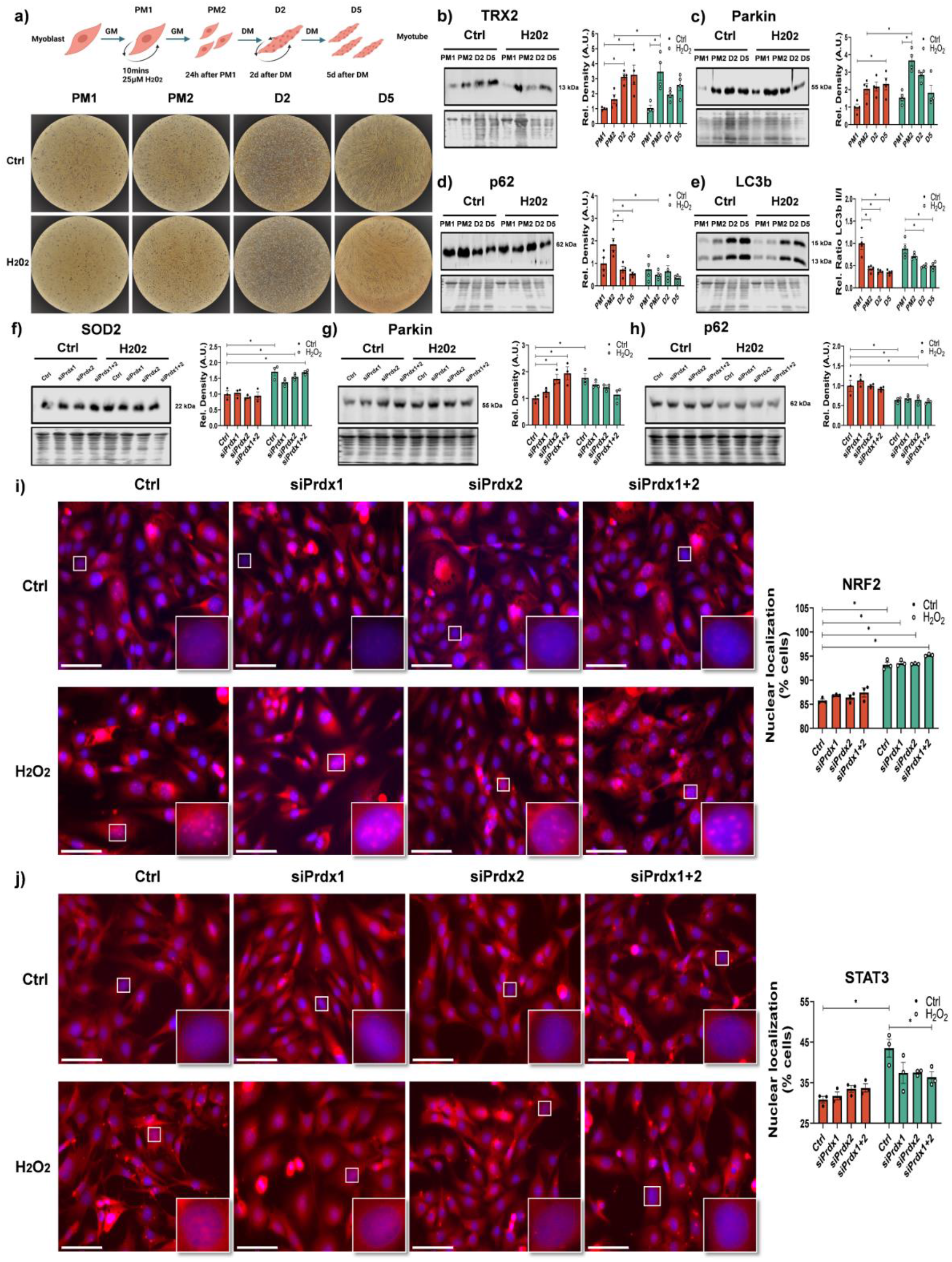
Schematic of approach for proliferation and differentiation of C2C12 cells following 10 min with 25 μM H_2_O_2_ (a). Western blot analysis expression during the proliferation and differentiation of C2C12s controls and cells treated for 10 min with 25 μM H_2_O_2_ for TRX2, Parkin, p62 and LC3 II/I (b-e). Cells with siPrdx1 and/or Prdx2 and treated with H_2_O_2_ and Western blot analysis for SOD2, p62 and Parkin (f-h). C2C12 cells with nuclear localisation of NRF2 and STAT3 following siPrdx1 and/or Prdx2 and treatment with H_2_O_2_, scale bar =75μm. Graphs are the mean +/-SEM and all experiments with n=3, two-way ANOVA analysis was used between groups and *p-*value of <0.05 was considered as statistically significant *(*p*<0.05). (a-k). *p values (b: Ctrl: PM1 vs Ctrl: D5= 0*.*0050, Ctrl: PM2 vs H2O2: PM2= 0*.*0330; c: Ctrl: PM1 vs Ctrl: D5= 0*.*0470, Ctrl: PM2 vs H2O2: PM2= 0*.*0084; d: Ctrl: PM2 vs Ctrl: D5= 0*.*0017, Ctrl: PM2 vs H2O2: PM2= 0*.*0011; e: Ctrl: PM1 vs Ctrl: D5< 0*.*0001; f: Ctrl: Ctrl vs H2O2: Ctrl= 0*.*0005; g: Ctrl: Ctrl vs Ctrl: siPrdx2= 0*.*0453, Ctrl: Ctrl vs H2O2: Ctrl= 0*.*0314; h: Ctrl: Ctrl vs H2O2: Ctrl= 0*.*0370; i: Ctrl: Ctrl vs H2O2: Ctrl< 0*.*0001; j: Ctrl: Ctrl vs H2O2: Ctrl= 0*.*0003)*.

**Supplementary Figure 3.**
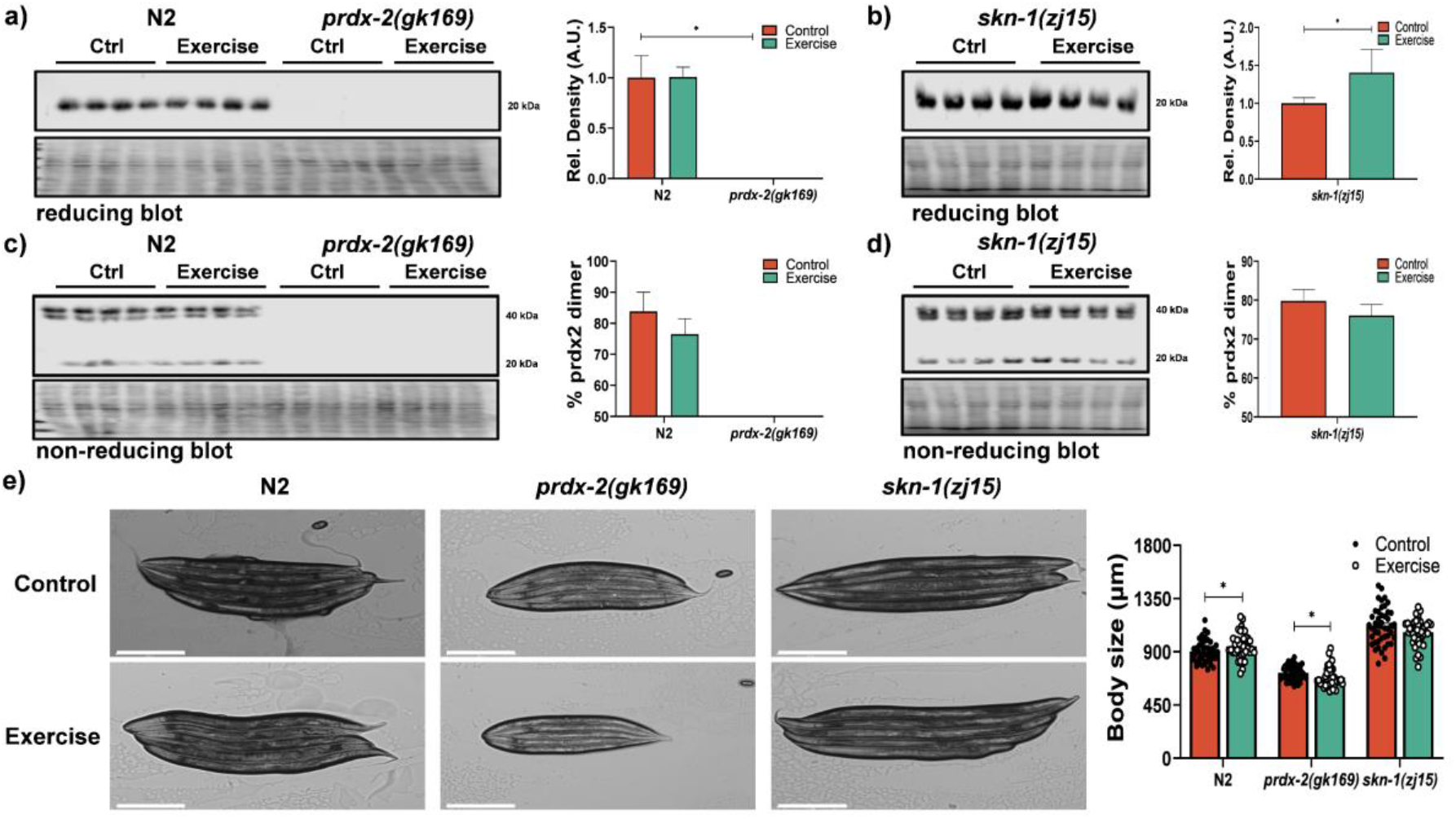
Immunoblotting for PRDX-2 expression in N2, *prdx-2* (a) and *skn-1* (b) strains following a 5day exercise protocol. Non-reducing immunoblot for analysis of monomer/dimer formation of PRDX-2 in N2, *prdx-2* (c) and *skn-1* (d) strains following 5 day exercise protocol. Body size of N2, *prdx-2 (gk163)* and *skn-1 (zj15)* strains, experiments were performed with at least 45 worms per experiment (e). Graphs are the mean +/-SEM, all experiments were performed n=3-4 and analysed by Student *t* test and *p-*value of <0.05 was considered as statistically significant *(*p*<0.05). *p values (a: N2: Control vs prdx-2 (gk163): Control< 0*.*0001; b: Control vs Exercise= 0*.*0420; e: Body size: N2: Control vs Exercise= 0*.*0340, prdx-2 (gk163): Control vs Exercise= 0*.*0107, skn-1 (zj15): Control vs Exercise= 0*.*0612)*.

**Supplementary Figure 4.**
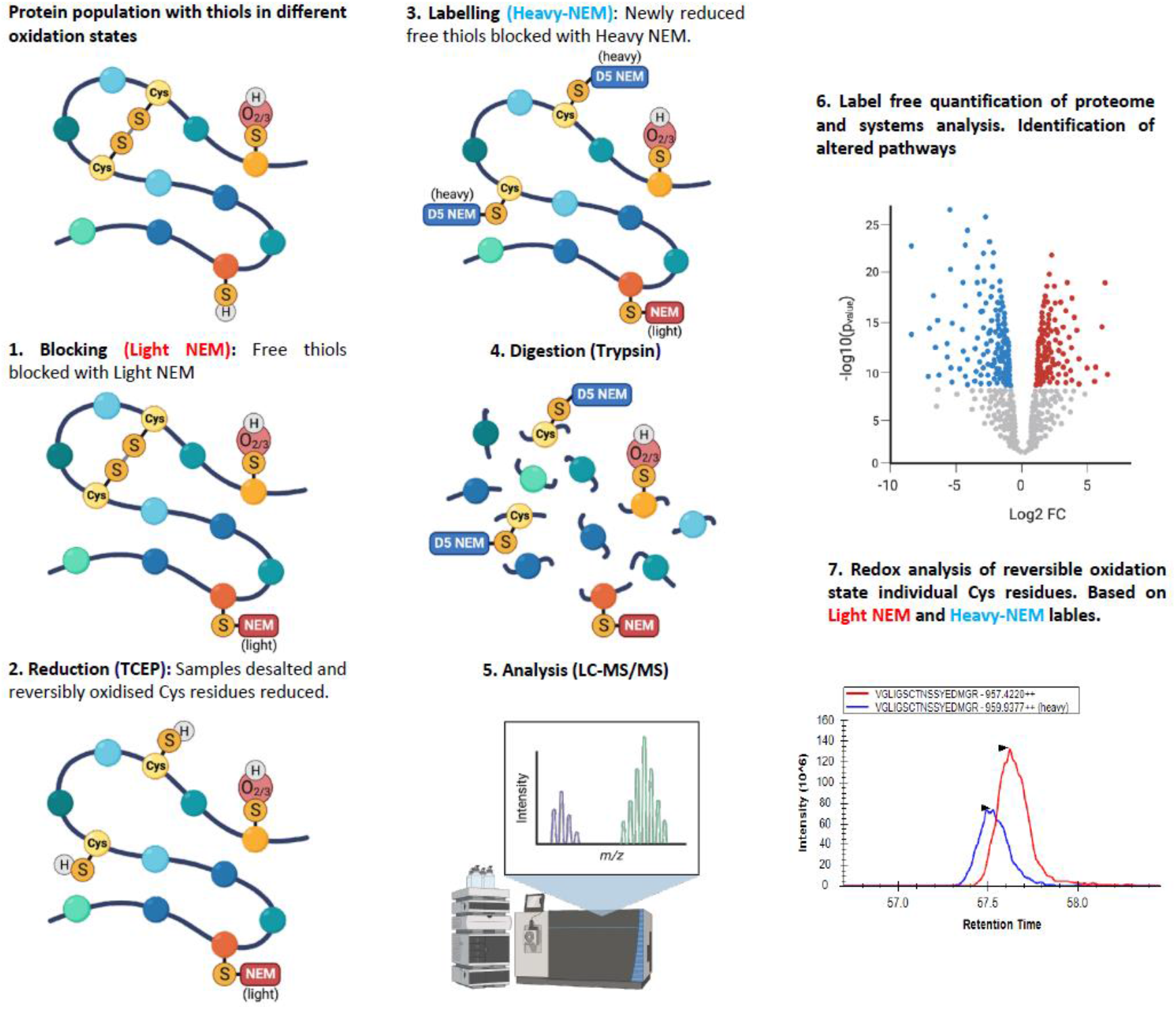
Schematic diagram of the redox proteomic approach to quantify protein abundance by label free quantification and relative quantification of the reversible oxidation state of peptides containing redox sensitive Cys residues.

**Supplementary Table 1.**
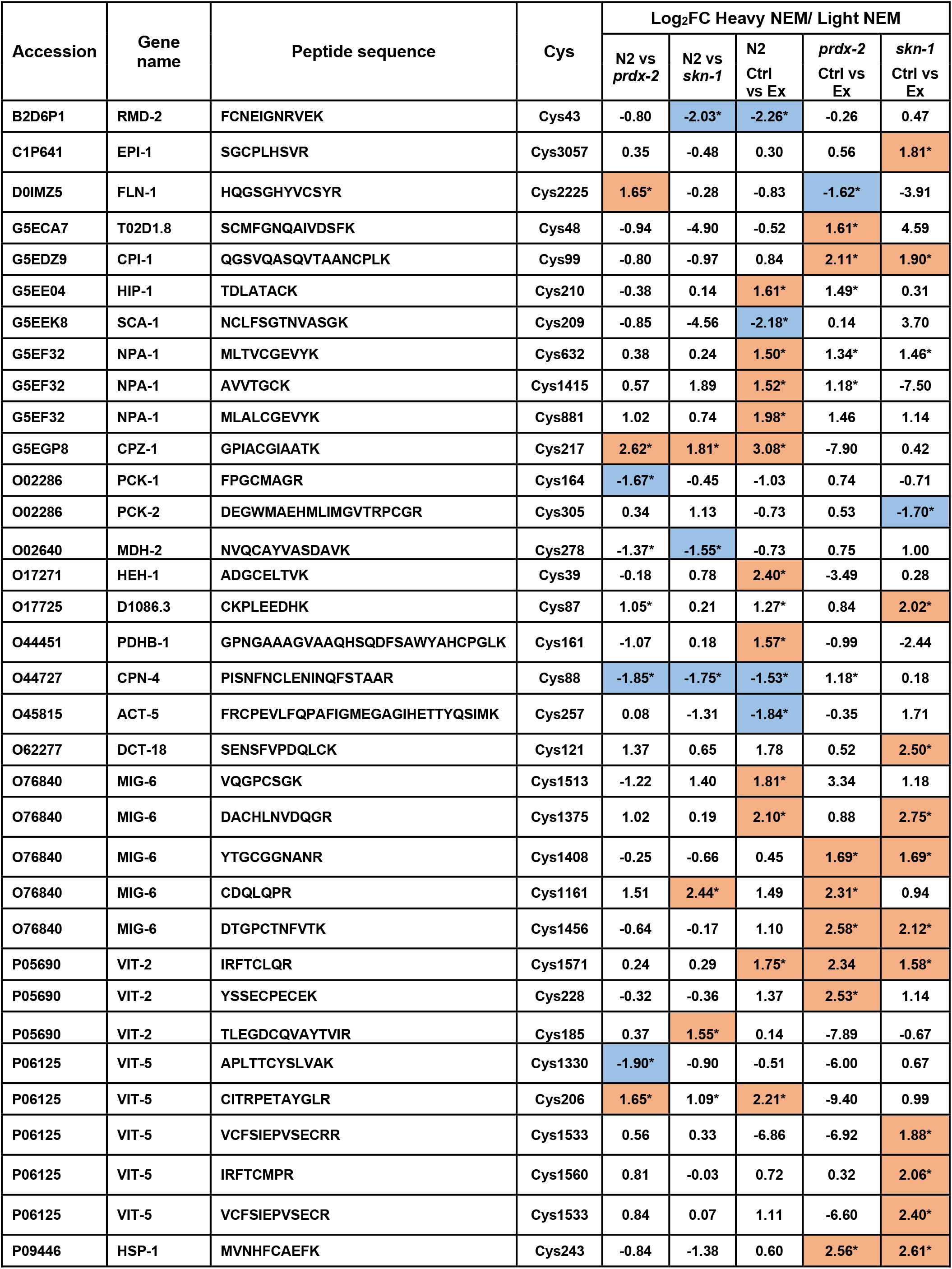
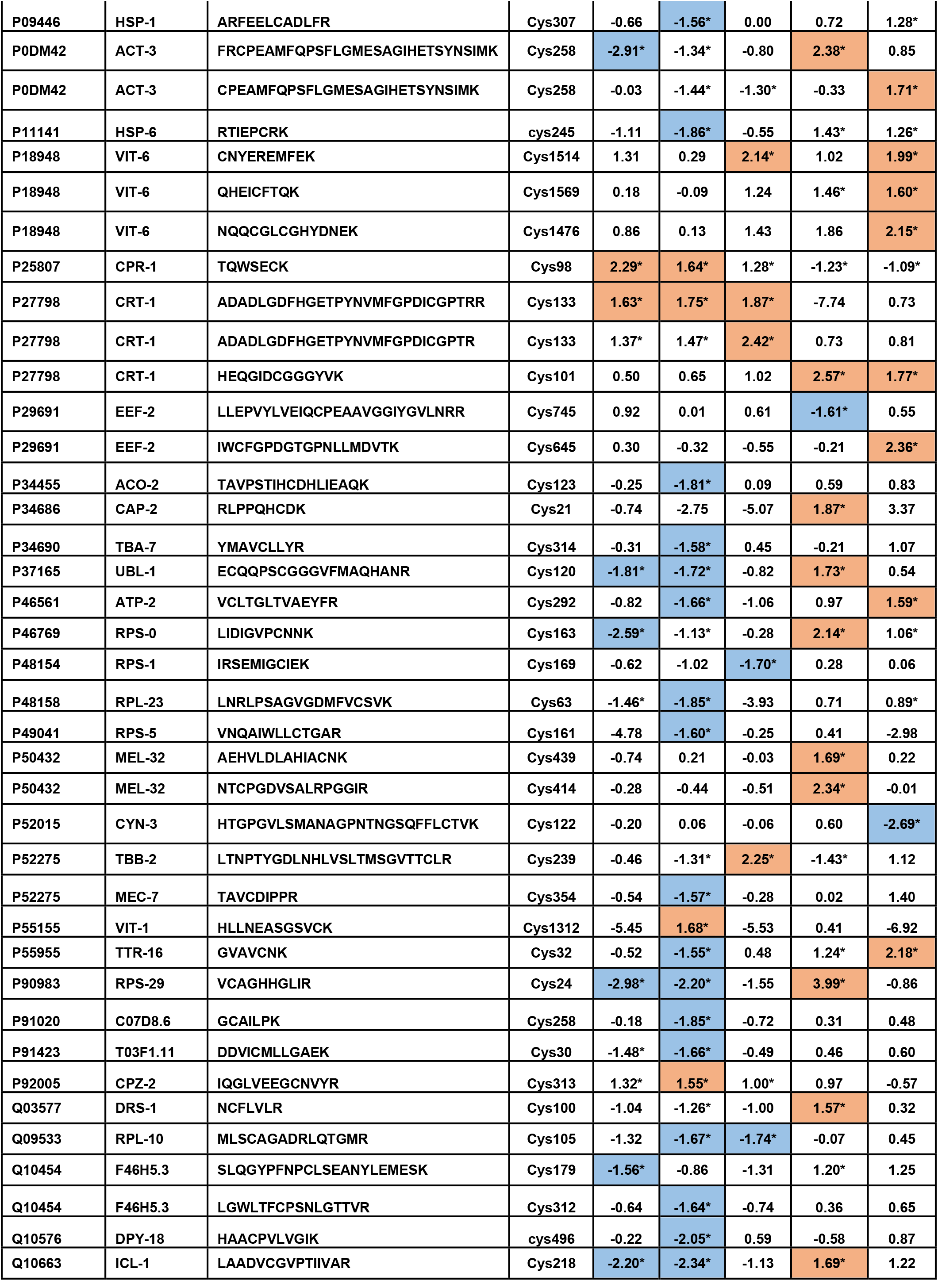
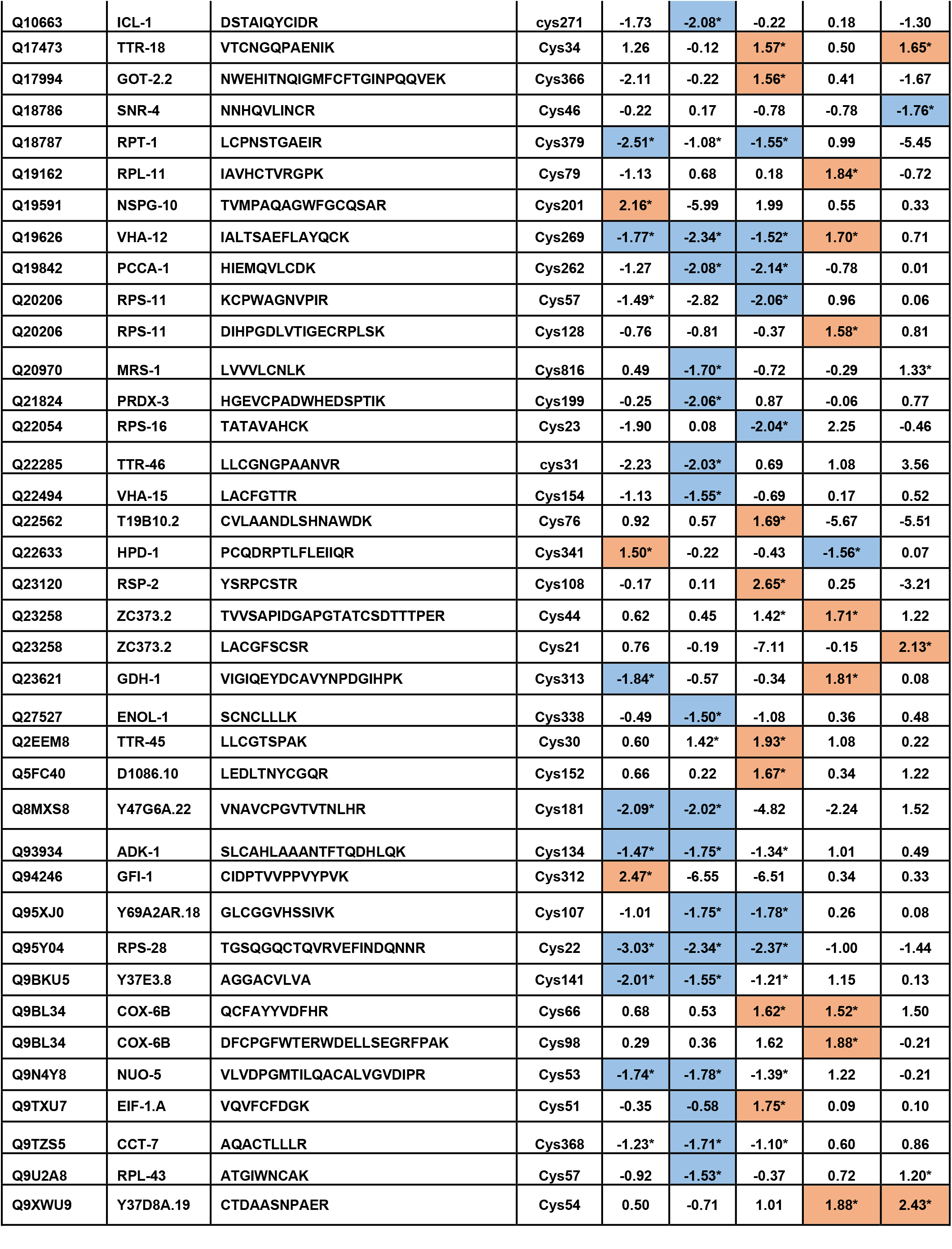
List of Cys containing peptides of where the redox state is significantly changed comparing the Log_2_Fold change of ratio of Heavy NEM:NEM labelling of Cys residues in non-exercised strains N2 Vs *prdx-2* and N2 Vs *skn-1* and following exercise compared to non-exercised controls in N2, *prdx-2* and *skn-1* strains. Peptides coloured blue are Cys residues more reduced and those in orange are more oxidised, * have p-value <0.05.

**Supplementary File 1**. Label free quantification proteomic data from N2, *prdx-2, skn-1* and *bli-3* strains with and without swimming exercise including differential analysis of proteins.

**Supplementary File 2**. Redox proteomic analysis of reversible oxidation state of Cys containing peptides labelled with both light and heavy NEM, Log_2_FoldChange and p-value.

